# Piezo1 couples fluid shear stress to adaptive genome dynamics by integrating cytoplasmic-nuclear mechanotransduction

**DOI:** 10.64898/2026.06.15.732233

**Authors:** Haymar Wint, Matthew S Graus, Yew Y Wong, Li-Kun Phng, Mathias Francois

## Abstract

Fluid shear stress (FSS) regulates endothelial morphology and function through flow-responsive gene expression programs. Yet, how mechanical forces are transmitted across cytoplasmic and nuclear compartments to regulate genome adaptive response remains unclear. Here, we show that FSS induces rapid nuclear remodeling characterized by nuclear compaction and apical nuclear indentations. These changes are driven by reorganization of perinuclear actin and microtubule cytoskeleton into apical linear cytoskeletal cables that constrain the nuclear surface. Concurrently, the mechanosensitive ion channel Piezo1 redistributes from the plasma membrane to these perinuclear deformations. Quantitative molecular imaging under flow reveals a transient adaptive cell state characterized by chromatin reorganization and epigenetic remodeling, accompanied by altered mobility of the flow-responsive transcription factor KLF2. Pharmacological inhibition of Piezo1 abolishes FSS-induced nuclear deformation and uncouples chromatin reorganization from KLF2 dynamic changes. Together, these findings reveal that endothelial mechanotransduction exploits physical principles of nuclear organization to regulate transcription factor behavior and adaptive genome responses across biological scales.

## Introduction

Blood vessels are lined by a monolayer of endothelial cells (ECs) that are continuously exposed to hemodynamic forces generated by circulating blood. Among these forces, fluid shear stress (FSS) is a dominant regulator of endothelial cell state and behavior, essential to preserve vessel integrity. In response to laminar FSS, ECs align in the direction of flow and adopt an anti-inflammatory, atheroprotective phenotype whereas exposure to disturbed or oscillatory flow promotes pro-inflammatory and pro-atherogenic responses vascular homeostasis [1–3]. These adaptive responses are mediated by mechanotransduction pathways that convert mechanical cues into biochemical signals to maintain vascular homeostasis.

Mechanotransduction in ECs has classically been attributed to signaling at the plasma membrane involving mechanosensitive ion channels, adhesion molecules, membrane receptors, and the cytoskeleton [1, 4–6] adaptations through epigenetic mechanisms, including chromatin remodeling and shear-dependent histone modifications, which differ between atheroprotective [7, 8]. However, precise mechanisms by which mechanical forces directly shape chromatin organization and transcriptional outputs remain elusive.

Emerging evidence indicates that mechanotransduction is not confined to the plasma membrane but extends to the nucleus, which functions as a mechanosensitive organelle. Nuclear morphology, stiffness, and chromatin organization are dynamically regulated by cytoskeletal forces transmitted through the LINC complex and nuclear lamina components, including Lamin A/C [9–11]. Importantly, cytoskeletal networks that transmit forces to the nucleus are themselves primary downstream effectors of endothelial shear stress sensing, positioning the nucleus as an active participant in endothelial mechano-adaptation.

Piezo1 is a well-established mechanosensitive ion channel that mediates endothelial adaptation to FSS by triggering calcium influx and proteolytic cleavage of actin cytoskeletal and focal adhesion proteins [4]. Through these effects, Piezo1-dependent signaling directly reshapes cytoskeletal architecture, providing a potential mechanical link for force transmission from the plasma membrane to the nucleus. Moreover, Piezo1 and PECAM1 interact at cell-cell junctions in endothelial force sensing [12], further positioning Piezo1 within junctional–cytoskeletal networks capable of relaying shear-dependent forces. Beyond its role in cytoskeletal remodeling, Piezo1 regulates flow-responsive transcriptional programs, including induction of the expression of the transcription factor KLF2 during heart valve development of zebrafish and in ECs [13, 14]. Further, MyoD family-inhibitor containing domain (MDFIC) transcriptional regulator has been shown to act as a Piezo sub-unit in lymphatic endothelial cells [15]. These findings suggest that Piezo1 signaling converges on transcriptional regulation through both biochemical and mechanically mediated pathways.

Consistent with a role in nuclear mechanoregulation, recent work demonstrates that Piezo1 can modulate nuclear mechanical properties through calcium-dependent actin remodeling, as revealed by its pharmacological activation with the Yoda1 agonist [16]. Together, these findings suggest a model in which Piezo1-dependent mechanosensing synergise cytoskeletal force with nucleus dynamics, thereby influencing downstream chromatin organization and genome effectors activity. However, how Piezo1 integrates cytoskeletal and nuclear mechanotransduction to instruct genome function under fluid shear stress remains unclear.

In this study, we characterize a Piezo1-associated mechanical axis that links endothelial shear sensing to adaptive nuclear and genome regulation under FSS. In addition to its established role as a plasma membrane mechanosensor, we observe Piezo1 at cytoskeletal indentations along the nuclear envelope during FSS adaptation. By combining quantitative nuclear imaging, chromatin texture analysis, super-resolution microscopy, and single-molecule tracking, we investigate how flow-induced cytoskeletal remodelling is connected to nuclear envelope adaptation, chromatin reorganization, and transcription factor dynamics under FSS. Together, this work establishes a multiscale framework for understanding how endothelial cells convert mechanical forces from flow into coordinated changes in nuclear architecture, genome organization, and flow-responsive transcriptional regulation.

## Results

### Fluid shear stress drives coordinated perinuclear cytoskeletal remodeling and nuclear deformation in endothelial cells

As previously established, prolonged laminar fluid shear stress (2 Pa FSS) induced the alignment of human aortic endothelial cells (HAECs), transitioning from cobblestone morphology under static conditions to an elongated shape after 24 and 48 hr FSS exposure (Supp fig.1A). This cellular alignment was accompanied by a coordinated reorganization of cytoskeletal components, including actin and microtubules (Fig 1A), which were required for endothelial adaptation to shear stress. Notably, as early as 6 hr after the onset of FSS, actin and microtubule cytoskeleton formed prominent cables spanning the apical surface of the nucleus (Fig 1B, Supp Fig 1B), in contrast to static HAECs that exhibit a mesh-like perinuclear cytoskeletal cage on the apical region of the nucleus. Closer examination of nuclear morphology revealed that ECs under static conditions exhibited nuclei with smooth and uniform nuclear membranes, whereas cells exposed to flow displayed pronounced wrinkling of the nuclear membrane as indicated by a stripe of low DAPI signal intensity bordered by intense Lamin A/C staining (white arrowheads in Fig. 1B and Supp Fig. 1C). These deformations appeared as indentations on the apical nuclear surface in orthogonal views (arrows in Fig. 1B). High-resolution imaging further revealed that actin and microtubule bundles aligned within these nuclear indentations (arrowheads in Fig. 1B and D). Line intensity profile analysis (Fig.1C) revealed the overlapping of actin and microtubules at the site of nuclear indentation. The close spatial association between cytoskeletal bundles and the deformed nuclear envelope, together with the local reduction in DAPI signal, is consistent with cytoskeletal impingement contributing to indentation formation. Quantification confirmed a marked increase in the proportion of cells exhibiting nuclear indentations after 6 hr of FSS exposure, whereas such features were infrequent under static conditions (Fig. 1C). These observations reveal early remodelling of the perinuclear cytoskeleton and nuclear membrane upon exposure to FSS.

**Fig 1.**
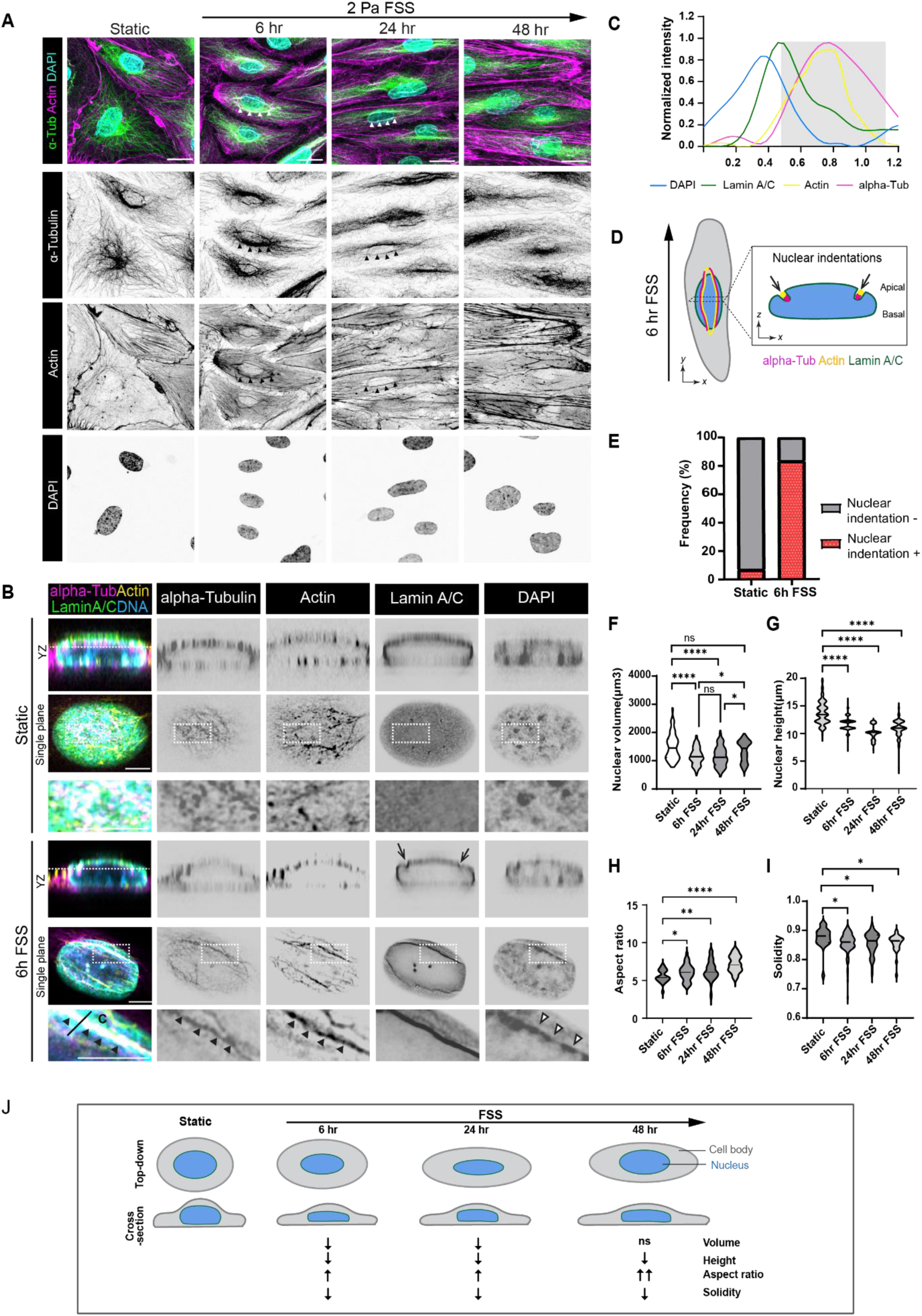
Fluid shear stress induces temporal cytoskeletal remodeling and nuclear morphological changes in HAECs. (A) Representative maximum intensity projections (MIPs) of HAECs cultured under static condition or exposed to laminar fluid shear stress (FSS, 2 Pa) for 6, 24 and 48 hrs. Cells were stained for α-Tubulin (green), F-actin (magenta) and nuclei (DAPI, cyan). Scale bar, 20μm. (B) Representative Airyscan images showing orthogonal YZ views and apical single-plane sections of HAECs under static conditions or exposed to 6 hr FSS. HAECs were stained for α-Tubulin (magenta), F-actin (yellow), Lamin A/C (green) and nuclei (DAPI, blue). Arrows, apical nuclear indentations. Enlarged images show regions highlighted by dashed boxes. Black arrowheads, apical linear actin– and microtubule-positive cables within nuclear indentations. White arrowheads, stripe of low DAPI signal on apical surface of nucleus. Scale bar, 5 μm. (C) Fluorescence intensity profile of Lamin A/C (green), α-Tubulin (magenta), actin (yellow) and DAPI (blue) along the indicated line scan across the nuclear indentation region under FSS conditions in (B), demonstrating enrichment of α-Tubulin and actin within nuclear indentation (grey box shows lower intensity signals of Lamin A/C and DAPI in nuclear indentation region). (D) Schematic illustration showing side view of the cell under FSS, highlighting apical nuclear indentation relative to the basal surface. Coloured outlines represent the nuclear membrane. (E) Percentage of HAECs displaying at least one apical nuclear indentation under static or 6 hr FSS conditions, n=100 cells from three independent experiments. (F–I) Quantitative analysis of nuclear volume (E), height (F), aspect ratio (G) and solidity (H) in HAECs exposed to 2 Pa FSS for the indicated durations. n=150 cells from three independent experiments). Data are presented as violin plots with median and quartile distribution. Statistical significance was determined by Welch’s T-test. Asterisks denote statistical significance (**** p < 0.0001, ***p < 0.001, **p <0.01, *p < 0.05, ns = no significant). (J) Working model summarizing endothelial nuclear remodeling during adaptation to FSS. Flow exposure induces endothelial elongation resulting in progressive nuclear deformation characterized by nuclear flattening, reduced nuclear volume, solidity and increased nuclear elongation over time.

We next investigated whether nuclear indentation was associated with measurable changes in nuclear morphology across a FSS exposure time course. Nuclear morphometric analysis revealed that 48 hr of FSS exposure reduced nuclear volume (Fig.1F) and height (Fig.1G) while increasing nuclear aspect ratio (Fig.1H). These changes occur as early as 6h of FSS exposure, indicating sustained nuclear flattening and elongation over time. Moreover, nuclear solidity (defined as the ratio of measured nuclear volume to the volume of its convex hull) progressively decreased following FSS exposure (Fig. 1I), reflecting the development of irregular nuclear contours and surface indentations. Notably, this reduction coincided with the formation of apical linear actin–microtubule bundles and nuclear indentations. Together, these findings reveal that shear stress-induced cytoskeletal remodeling is coupled with time-dependent progressive deformation of the apical nuclear surface and reshaping of nuclear morphology in ECs (Fig. 1J). This observation prompted us to survey flow-induced global nuclear changes from nuclear architecture to single molecule behavior of genome effectors (Fig. 2A).

**Fig 2.**
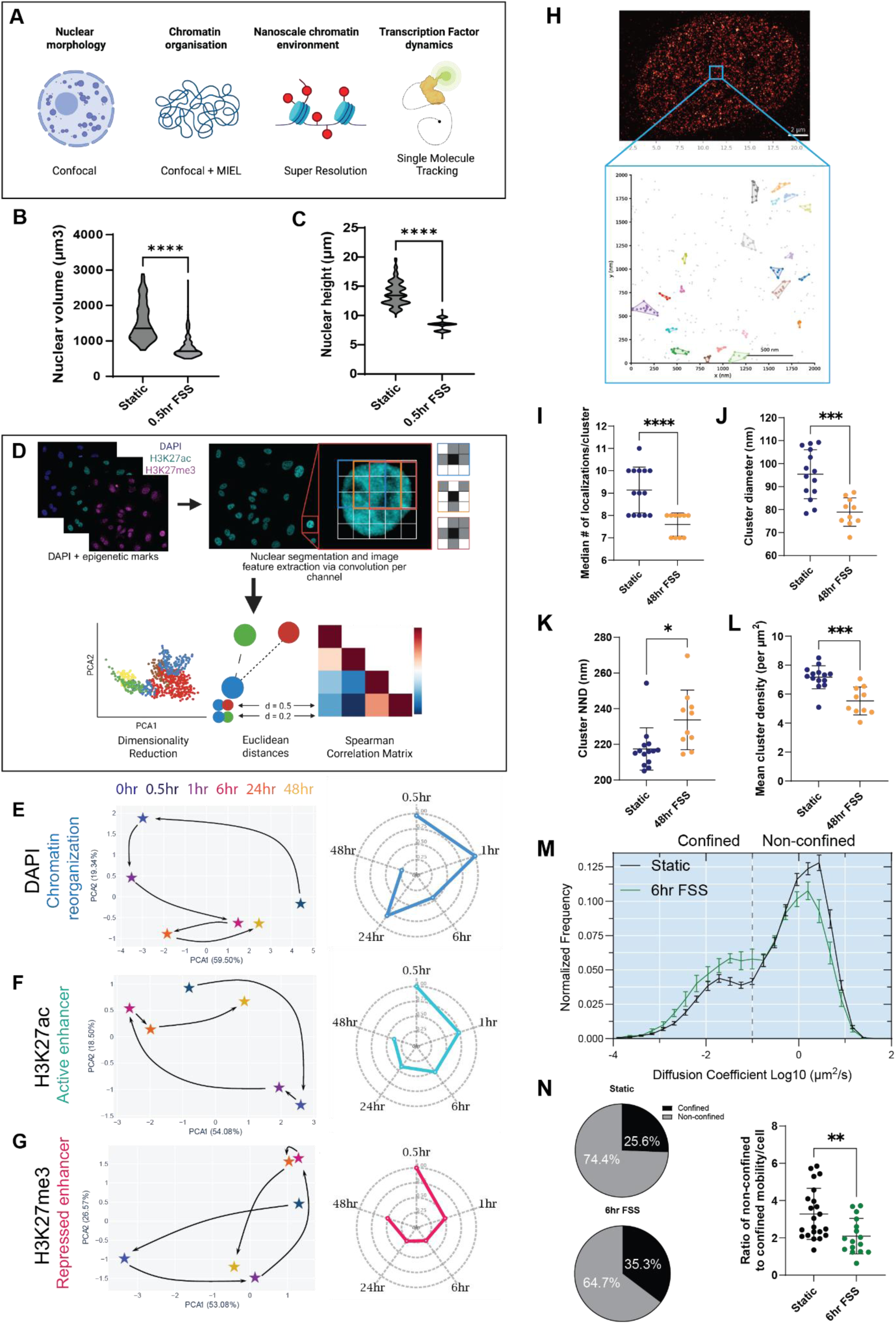
Fluid shear stress drives early remodeling of nuclear morphology, genome organization, and chromatin organization. (A) Schematic overview of the multiscale imaging and analysis pipeline used to characterize nuclear and chromatin responses to FSS. (B–C) Nuclear morphology of HAECs after 0.5 hr exposure to 2 Pa FSS. (B) Nuclear volume and (C) nuclear height. Data are violin plots with median and interquartile distribution; n = 150 cells from three independent experiments. Statistical significance determined by t-test, ****p < 0.0001. (D) Schematic of the MIEL analysis pipeline. Confocal images were acquired per condition and separated by fluorescence channel. Nuclei were individually segmented using DAPI as a mask, and 252 texture– and edge-based features were extracted from pixel intensity patterns within each segmented nucleus, capturing spatial variation in fluorescent signal distribution relative to neighboring pixels. Feature variation across the nuclear population was summarized by PCA, and pairwise relationships between conditions were quantified using Euclidean distance and Spearman correlation. (E–G) MIEL analysis across the FSS time course for (E) DAPI, (F) H3K27ac and (G) H3K27me3. PCA plots show PC1–PC2 feature space; stars indicate condition centroids and arrows indicate temporal trajectory. Radar plots display Euclidean distance of each timepoint centroid relative to 0 hr, normalized independently for each channel. n ≥ 750 nuclei per condition from three biological replicates.PERMANOVA was performed using retained PCs explaining >80% of variance: DAPI, p = 0.002; H3K27ac, p = 0.091; H3K27me3, p = 0.617. (H) Representative H3K27ac dSTORM localization map of a HAEC nucleus. Inset shows a 2 µm × 2 µm ROI with DBSCAN-identified H3K27ac clusters; unclustered localizations are shown in grey. Scale bar, 2 µm. (I–L) dSTORM quantification of H3K27ac nanocluster characteristics under static conditions and after 48 hr FSS. Each point represents one cell unless otherwise indicated; static n = 14 cells and 48 hr FSS n = 10 cells from two biological replicates. Data are mean ± SD; Welch’s t-test. (I) Nanocluster density, (J) equivalent diameter, (K) localizations per nanocluster and (L) whole-nucleus nearest-neighbor distance. (M) KLF2 mobility distribution in HAECs exposed to static conditions or 2 Pa FSS for 6 hr. Histogram shows normalized KLF2 diffusion coefficients; dotted line indicates the threshold between confined and non-confined mobility states. Data represent pooled trajectories from n ≥ 16 cells per condition across three biological replicates; error bars represent SEM. (N) Quantification of KLF2 mobility states. Pie charts show confined and non-confined populations; dot plot shows the non-confined/confined ratio per cell. Data are mean ± SD; Welch’s t-test, **p = 0.0032.

### FSS elicits a coordinated adaptive response of both nuclear architecture and genome organization across scales

Because mechanotransduction operates on a timescale of seconds to minutes, rapid nuclear and genome-associated changes may represent early drivers of endothelial adaptation to flow. However, how nuclear morphology variations and chromatin reorganization unfold during the initial phases of the endothelial response to FSS remains unclear. We therefore designed a multimodal, multiscale imaging strategy to examine FSS-induced nuclear and genome remodelling across time and space, spanning nuclear morphology, mesoscale chromatin organization, H3K27ac nanoscale environment, and transcription factor (KLF2) dynamics at single-molecule resolution (Fig. 2A).

We exposed HAECs to FSS for 0.5hr and quantified changes in nuclear morphology. We observed that FSS rapidly significantly reduced nuclear volume and height compared with static conditions (Fig. 2B–C). This result prompted us to ask whether abrupt changes in nuclear morphology are accompanied by coordinated remodelling of chromatin organization, epigenetic mark distribution, and transcription factor mobility over time.

We first performed microscopic imaging of epigenetic landscapes (MIEL, [20]) on HAECs exposed to FSS for 0.5 hr, 1 hr, 6 hr, 24 hr, and 48 hr to determine whether early nuclear remodelling coincides with broader changes in chromatin organization (Fig. 2D–G). MIEL quantifies fluorescent signal and texture features from individual segmented nuclei (Fig. 2D) to assess chromatin reorganization [17]. DAPI was used to assess global chromatin origami, while H3K27ac and H3K27me3 were used to examine active and repressive epigenetic environments, respectively. Because each marker reflects a distinct aspect of genome organization, each channel was analyzed independently (Fig. 2E–G; Supp. Fig. 2B–J).

PERMANOVA confirmed that DAPI texture features were significantly reorganized by FSS, while the epigenetic mark channels showed trends that did not reach significance. The PCA and radar plots reveal a consistent directional trajectory across all three independently analyzed channels. Although the strength of the response differed by channel, all three markers showed a shared temporal pattern, with the greatest displacement from baseline occurring during the early 0.5–1 hr window, followed by a progressive partial reversion toward the 0 hr state (radar plots in Fig. 2E–G). Euclidean distance and Spearman correlation analyses support these trajectories (Supp. Fig. 2E-J). DAPI features showed the strongest multivariate separation, indicating that global chromatin organization is an early and prominent nuclear response to FSS. In contrast, H3K27ac and H3K27me3 followed distinct return trajectories: H3K27ac showed early displacement followed by a trend back toward baseline by 48 hr, whereas H3K27me3 partially returned toward baseline at 6–24hr before increasing again by 48 hr. These findings suggest that active and repressive chromatin environments respond to FSS with distinct temporal dynamics, consistent with rapid force transmission to chromatin through the LINC complex and slower histone-mark redistribution through chromatin-modifying enzymes [7, 21, 22]. Together, these results support a two-step model in which an early adaptive genome response is followed by a maintenance phase that approaches, but does not fully recapitulate, the static state.

### FSS remodels nanoscale H3K27ac organization

Although H3K27ac texture features trended back toward baseline at the MIEL scale by 48 hr, MIEL cannot determine whether nanoscale H3K27ac organization is similarly restored. We therefore used direct stochastic optical reconstruction microscopy (dSTORM) to compare H3K27ac nanoscale organization (∼20–30 nm) under static conditions and after 48 hr of FSS. H3K27ac was selected because it marks active enhancer elements and transcriptionally permissive chromatin states (Fig. 2H).

dSTORM analysis revealed that FSS caused H3K27ac texture changes detected by MIEL are associated with two related nanoscale changes, 1) individual H3K27ac nanoclusters are less populated (Fig. 2I) smaller (Fig. 2J, Supp. Fig. 2K), and less round (Supp. Fig. 2M). In addition, the remaining nanoclusters are distributed differently relative to one another (Fig. 2K-L, Supp Fig 2M). This is further supported by Ripley’s L analysis of nanocluster distribution showing an altered cluster-to-cluster organization across nanoscale distance ranges (Supp. Fig. 2N-O).

### FSS shifts KLF2 toward a chromatin-associated mobility state

Having established that FSS remodels nuclear architecture and active chromatin organization across multiple scales, we next asked whether these structural changes are accompanied by altered dynamics of a flow-responsive transcriptional regulator. We focused on Krüppel-like factor 2 (KLF2), a well-established flow-responsive transcription factor in ECs [18, 19], and performed single-molecule tracking (SMT) of Halo-tagged KLF2 in live cells under static conditions and after 6hr of FSS.

SMT revealed the population of KLF2 molecules significantly increased its confined mobility (e.g chromatin-associated ¥ state) after 6 hr of FSS (Fig. 2M–N). This indirectly reflects increased chromatin engagement, which is conditioned by change in accessibility, nuclear crowding, or a combination of these mechanisms.

Altogether, this combination of quantitative molecular imaging approaches indicates that FSS drives a temporally ordered remodelling across multiple structural and functional scales. Nuclear morphology and bulk chromatin organization change rapidly, consistent with early force transmission to the nucleus, before chromatin organization partially returns toward a flow-based equilibrium close to a static state. In contrast, super-resolution imaging unveiled persistent nanoscale remodelling of active enhancers, suggesting that active chromatin retains a longer-lived mechanical imprint of the FSS state. The accompanying increase in KLF2 confinement further links these structural changes to a functional adaptation of transcription factor behavior during endothelial response to flow. These results raised the possibility that perinuclear cytoskeletal remodelling acts as a key integrator of adaptive genome responses to flow. We therefore tested whether disrupting cytoskeletal remodelling alters the transmission of mechanical information and therefore impairs nucleus and genome adaptive response.

### Impaired cytoskeletal remodeling disrupts flow-induced nuclear architecture and chromatin organization

Previous studies have shown that cytoskeletal organization influences nuclear spatial architecture and chromatin dynamics in Drosophila S2R+ cells [20], prompting us to examine whether FSS-induced changes in nuclear morphology and chromatin organization are mediated by cytoskeletal reorganization. Nocodozole, taxol and latrunculin B (Lat. B) treatment significantly reduced FSS-induced nuclear indentations by ∼100%, ∼50% and ∼20%, respectively (Fig. 3A-B). As shown in Fig.1F, 6 hr exposure to FSS resulted in a 20%, 15%, and ∼8% reduction in nuclear volume, height, and solidity when compared with static controls. However, perturbation of microtubule dynamics with the depolymerization agent, nocodazole, and the stabilizing agent, taxol, attenuated FSS-induced changes in nuclear morphology (Fig. 3A). When compared to control, both drugs significantly increased nuclear volume, height and solidity (nocodozole: 120∼%, ∼42% and ∼ 4%, respectively; taxol: 120%, 32% and ∼7%, respectively) and significantly decreased nuclear aspect ratio (nocadazole, 11.7%; taxol, 10.8%; Fig. 3E). Disruption of actin filaments with Lat. B resulted in a partial reduction of the FSS-induced reduction and producing a ∼22% increase relative to DMSO-treated cells. (Fig. 3C-F). Together, these observations highlight that dynamic remodeling of both microtubule and actin cytoskeleton plays a role in FSS-driven nuclear morphological changes.

**Figure 3.**
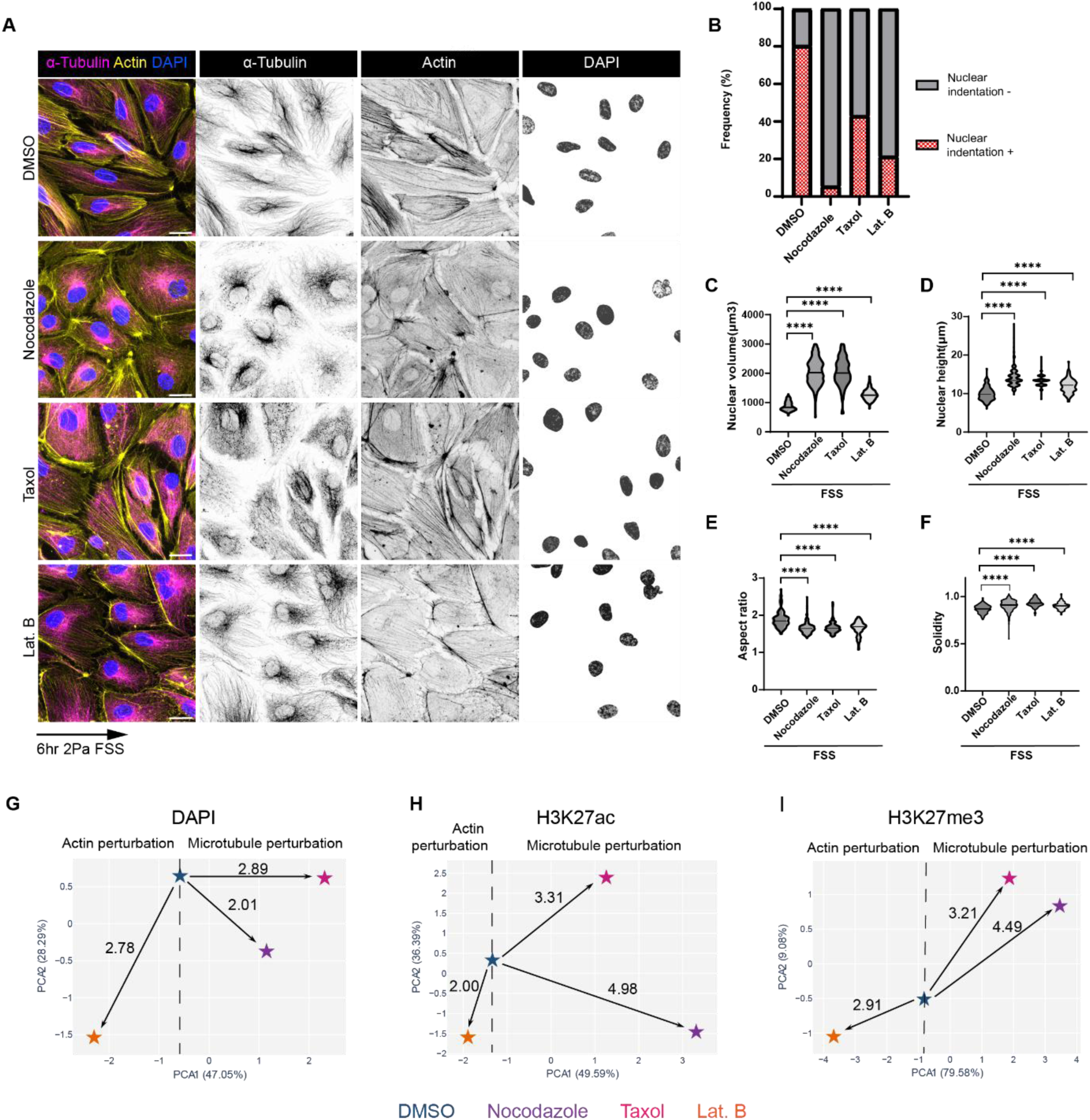
Cytoskeletal perturbation prevents FSS induced nuclear indentation formation and nuclear morphological changes in HAECs. (A) Representative MIPs of HAECs exposed to 2 Pa FSS for 6hrs in the presence of DMSO, nocodazole, taxol or latrunculin B (Lat. B). Cells were stained for α-Tubulin (magenta), actin (yellow), and DAPI (blue) channels. Scale bar, 20 μm. (B) Percentage of HAECs displaying at least one nuclear indentation under each condition. (n=100 from three independent experiments). (C–F) Quantitative analysis of HAEC nuclear volume (C), height (D), aspect ratio (E) and solidity (F) under FSS following cytoskeletal perturbation. n=150 for three independent experiments. Data are presented as violin plots with median and quartile distribution. Statistical significance was determined by Welch’s T-test. Asterisks denote statistical significance (**** p < 0.0001, ***p < 0.001, **p <0.01, *p < 0.05, ns = no significant). (F–H) PCA of features extracted from (F) DAPI, (G) H3K27ac, and (H) H3K27me3 staining in HAECs exposed to FSS for 6 hr in the presence of DMSO, nocodazole, taxol, or Lat. B. PCA plots show the first two principal components for visualization. Each star represents the condition centroid in PC1–PC2 space, with arrows indicating the Euclidean distance from the DMSO centroid to each drug condition. Numbers above arrows indicate Euclidean distance relative to the DMSO condition. The dashed vertical line separates actin perturbation conditions from microtubule perturbation conditions. PERMANOVA was performed using the retained multivariate PC space, explaining > 80% of total variance in each channel, and confirmed a significant shift in DAPI-derived features (p = 0.016) and H3K27ac-derived features (p = 0.005), but not H3K27me3-derived features (p = 0.303). Variance explained by PC1 and PC2 is indicated on each axis. Analysis was performed on ≥ 316 nuclei per condition across two biological replicates for drug-treated conditions and four biological replicates for the DMSO condition.

We next asked how perturbed cytoskeletal dynamics and accompanying loss of FSS-induced change in nuclear morphology affects the reorganization of chromatin. Actin and microtubule networks play a balancing act in [21, 22]. While actin increases tension within the network and aligns the actin cap, microtubules generate a compression-bearing network that resists acto-myosin tension and maintains cellular architecture. To test the role of these two cytoskeleton on chromatin organization in our system we performed MIEL analysis on cells exposed to 6 hr FSS in the presence of nocodazole, taxol or Lat. B (Fig. 3G-I, Supp. Fig. 3A-C), PERMANOVA confirmed that DAPI and H3K27ac texture features were significantly reorganized, while H3K27me3 showed a trend that did not reach significance. Despite independent analysis, all three channels shared a distinct trend where both microtubule and actin pharmacological disruption drove chromatin reorganization, but in opposing directions within PCA space (Fig. 3G–I). This suggests that actin and microtubule networks regulate chromatin organization through distinct mechanisms [21, 22], even though both drugs induce an increase in nuclear volume. This pattern is further supported by Euclidean distance and Spearman correlation analyses across all channels (Supp. Fig. 3D–I). Thus, FSS-induced nuclear remodeling does not impose a uniform chromatin outcome. Rather, actin– and microtubule-dependent mechanical inputs can produce similar changes in nuclear volume while driving divergent chromatin and epigenetic states.

### Fluid shear stress induces changes in Piezo sub-cellular distribution in HAECs

Having established that cytoskeletal remodeling governs the translation of mechanical cues into distinct chromatin states, we next sought to identify the upstream mechanosensor responsible for coordinating this multi-scale adaptive response. To address this, we examined the temporal redistribution of Piezo1 during exposure to fluid shear stress. Piezo proteins are mechanosensitive channels that allow Ca²⁺ influx and are activated in ECs in response to shear stress [4]. Piezo1 has also been implicated in shear stress-induced nuclear shrinkage in epithelial systems [23], where Piezo1-mediated Ca^2+^ influx was proposed to activate cytoskeletal contractility to compressive forces that physically reduce nuclear volume [21]. This is consistent with the established role of Ca²⁺ as a second messenger that rapidly reorganizes actomyosin and microtubule cytoskeletons. Further Ca²⁺ waves have been shown to traverse the nucleus driving the decompaction of heterochromatin [24]. These findings support a direct mechanotransductive pathway linking plasma membrane mechanosensing to cytoskeletal reorganization and subsequent changes in nuclear morphology and chromatin organization [16].

Under static conditions, Piezo1 is localized at cell-cell contacts centrosome and diffused throughout the cytoplasm (Supp Fig. 4A-B). Notably, FSS induced two distinct patterns of Piezo1 subcellular relocalization. After 6 hrs of FSS, Piezo1 became increasingly associated with actin and microtubule bundles spanning the apical nuclear surface (Supp. Fig. 4C). Live cell imaging of Piezo1-HaloTag further supported Piezo1 redistribution, revealing enhanced perinuclear accumulation after 6 hr FSS (Supp Fig. 5, Supplementary Video 1). After prolonged FSS exposure (24–48 hr), when ECs became fully aligned, Piezo1 showed polarized enrichment at the downstream end of the cell relative to flow direction (Supp. Fig. 4C & D). These observations reveal a dynamic spatiotemporal redistribution of Piezo1 in response to FSS and indicate a previously unrecognized spatial coupling between this mechano-receptor and nuclear remodeling during adaptation to flow.

**Figure 4.**
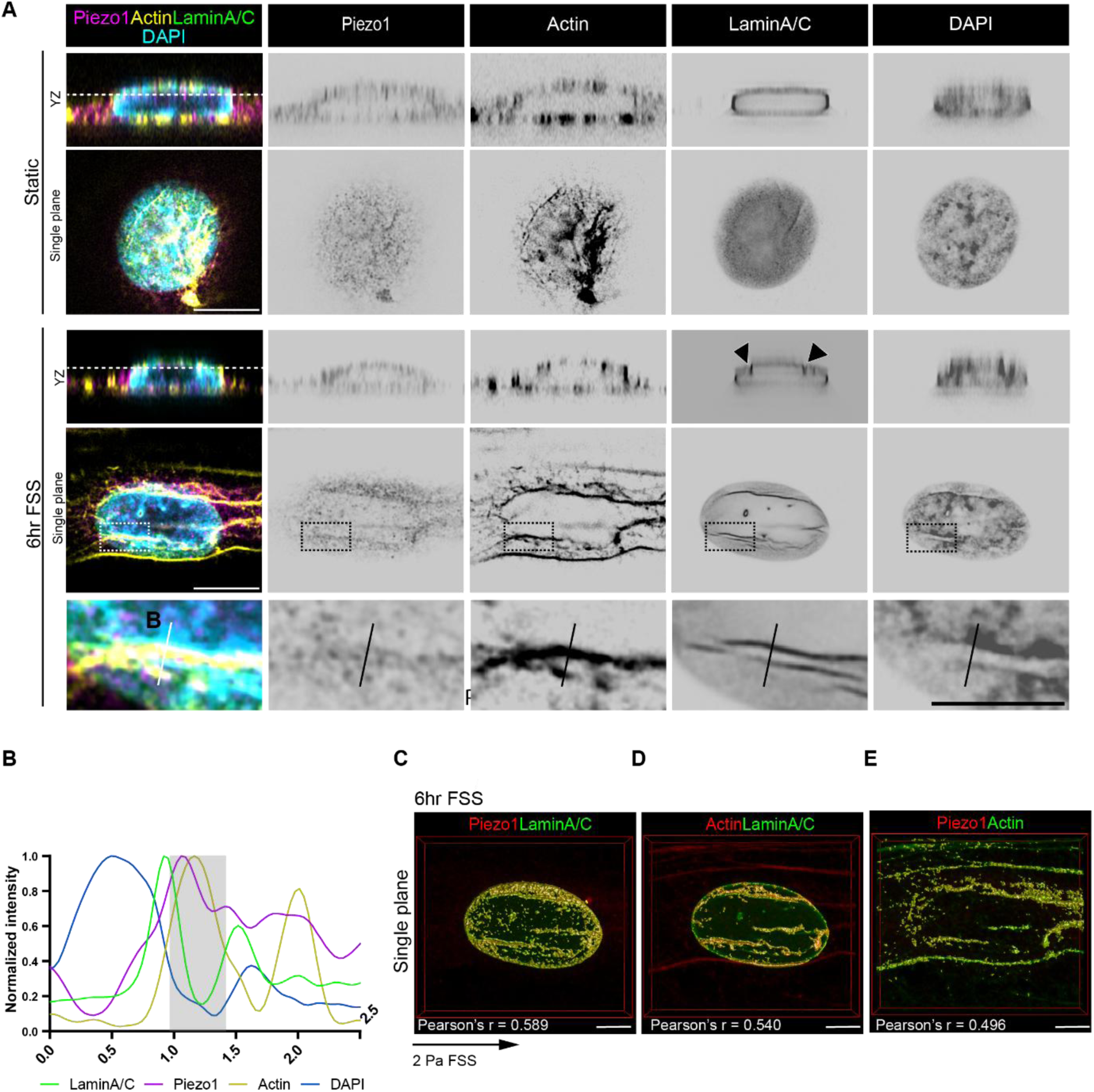
Fluid shear stress induces Piezo1 localization to apical nuclear indentations. (A) Representative Airyscan immunofluorescence images showing orthogonal YZ views and apical single-plane sections of HAEC cultured under static conditions or exposed to FSS for 6 hrs. HAECs were stained for Piezo1 (magenta), F-actin (yellow), Lamin A/C (green) and nuclei (DAPI, cyan). Arrowheads, apical nuclear indentations. Dashed box indicates localized enrichment of Piezo1 along linear actin cables that align within apical nuclear indentations shown at higher magnification below. Scale bars, 10 μm (upper panel) and 5 μm (lower panel). (B) Fluorescence intensity profile of Lamin A/C (green), Piezo1 (magenta), actin (yellow) and DAPI (blue) along the indicated line scan across the nuclear indentation region under FSS conditions in (A). demonstrating enrichment of Piezo1 and cortical actin within nuclear indentation (grey box, low LaminA/C levels). (C-E). Representative Airyscan images showing colocalization analysis of Piezo1 with Lamin A/C or Actin under 2 Pa FSS for 6 hr. Piezo1 is pseudocolored red (C and E); Lamin A/C in green (C and D); and Actin in red (D) and in green (E). Colocalization analysis was performed using Huygens Software, and Pearson’s correlation coefficients (r) are displayed in each panel. Merged signals indicating colocalization appear yellow. Scale bar, 5 μm.

**Figure 5.**
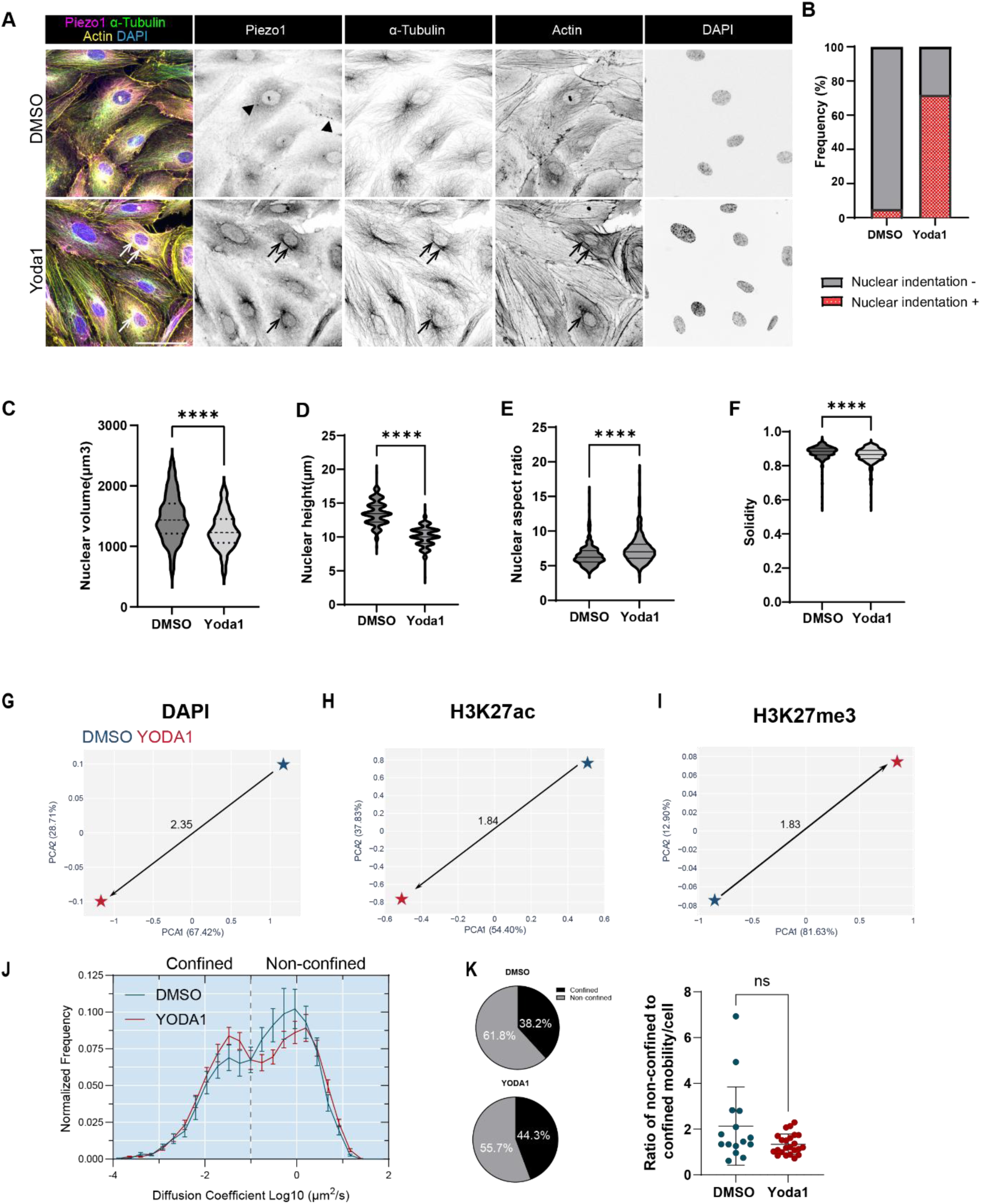
Activation of Piezo1 with Yoda1 phenocopies flow-induced nuclear indentation, nuclear remodeling, chromatin organization and TF mobility in endothelial cells. (A) Representative MIPs of HAECs cultured under static conditions and treated with DMSO or the Piezo1 agonist Yoda1. Cells were stained for Piezo1 (magenta), α-Tubulin (green), F-actin (yellow) and nuclei (DAPI, blue). Arrowheads indicate perinuclear localization of Piezo1 in DMSO treated cells. Arrows indicate Piezo1 localization at perinuclear linear cytoskeletal bundles after Yoda1 treatment. Scale bar, 50 μm. (B) Percentage of HAECs with at least one nuclear indentation in DMSO– and Yoda1-treated conditions, (n=100 cells from three independent experiments). (C-F) Quantitative analysis of nuclear morphology in HAECs treated with DMSO or Yoda1 under static conditions. Nuclear volume (C), height (D) aspect ratio (E) and solidity (F). (n=150 cells from three independent experiments). Data are presented as violin plots showing median and interquartile range. Statistical significance was determined by Welch’s T-test. Asterisks denote statistical significance (**** p < 0.0001). (G-I) PCA of features extracted from (G) DAPI, (H) H3K27ac, and (I) H3K27me3 staining in HAECs treated with DMSO (vehicle control) or Yoda1 (Piezo1 agonist). Each star represents the condition centroid in PC space with the Euclidean distance indicated. Variance explained by PC1 and PC2 is indicated on each axis. (J) Mobility distribution plot comparing diffusion coefficients in HAECs treated with DMSO or Yoda1. Dotted line indicates the threshold between confined and non-confined mobility states. Data presented as mean ± SEM. (K) Pie charts represent the proportion of the population found in either confined or non-confined state based on diffusion coefficient. Dot plot represents the ratio of non-confined to confined molecules per cell, from n ≥ 15 cells per condition from three biological replicates. Statistical significance was determined by Welch’s T-test, p = 0.096.

Since Piezo1 accumulated in the perinuclear region concomitantly with the appearance of nuclear indentations and perinuclear actin/microtubules bundles after 6 hr of FSS, we next examined the apical nuclear surface at higher resolution. Under static conditions, Piezo1 was distributed in a punctate pattern across the apical nuclear region (Fig. 4A). In contrast, after 6 hr of FSS exposure, Piezo1 adopted a linear distribution pattern that colocalized with apical linear actin bundles spanning the nucleus and impinging on the apical nuclear surface (boxed region in Fig. 4A). Line intensity profile analysis (Fig. 4B) revealed overlapping peaks of Piezo1 and actin signals within a nuclear indentation delineated by Lamin A/C staining and decreased DAPI staining intensity. The colocalization of Piezo1 with actin cytoskeleton and nuclear membrane is further supported by Pearson’s correlation coefficient analysis (Fig. 4C, Piezo1 and actin, r = 0.589, Piezo1 and Lamin A/C, r = 0.54 and actin and Lamin A/C, r = 0.534).

Together, these observations indicate that FSS induces the dynamic redistribution of Piezo1 along perinuclear cytoskeletal bundles that interface with the nucleus. The enrichment of Piezo1 at sites of nuclear indentation suggests that mechanosensing and nuclear remodeling may be spatially coordinated during endothelial adaptation to flow.

### Piezo1 activation is sufficient to partially phenocopy flow-induced nuclear and genome remodelling

While the dynamic redistribution of Piezo1 suggested a close association with cytoplasmic-nuclear remodeling under flow, it remained unclear whether Piezo1 activation is sufficient to drive the adaptive nuclear and genome responses induced by FSS. We therefore examined the consequences of pharmacological Piezo1 activation on nuclear morphology, chromatin organization, and transcription factor dynamics. We pharmacologically activated Piezo1 using its agonist Yoda1 under static conditions. Interestingly, Yoda1 treatment altered Piezo1 localization, resulting in its enrichment along perinuclear cytoskeletal bundles (arrows, Fig. 5A), mimicking the distribution observed in FSS-exposed cells. In contrast, DMSO-treated cells displayed a diffused cytoplasmic and perinuclear Piezo1 localization pattern (Fig. 5A). This redistribution suggested a potential association between Piezo1 activation, cytoskeletal reorganization, and nuclear remodeling. To assess whether this change in Piezo1 organization was accompanied by apical nuclear deformation, we quantified the proportion of nuclear indentation-positive cells. Under static conditions, Yoda1 markedly increased the proportion of nuclear indentation-positive cells (∼70%) compared with DMSO-treated controls (∼5%) (Fig. 5B), indicating that Piezo1 activation alone promotes nuclear indentation. Nuclear morphometric analysis further demonstrated that Yoda1 treatment phenocopied the effects of FSS by decreasing nuclear volume (Fig. 5C), height (Fig. 5D), and solidity (Fig. 5E) while increasing nuclear aspect ratio (Fig. 5F). Collectively, these observations suggest that Piezo1 activation is sufficient to engage a perinuclear cytoskeletal force-transmission pathway capable of remodeling nuclear morphology, although direct evidence that this response requires actin or microtubule bundles will require further perturbation experiments.

To uncover whether these structural changes extended to chromatin and epigenetic mark organization, we performed MIEL analysis across DAPI, H3K27ac, and H3K27me3 channels. Yoda1 treatment altered and chromatin-associated feature spaces compared with DMSO, with Yoda1-treated cells showing measurable separation from controls across all three imaging channels (Fig. 5G–I, Supp. Fig. 6A–F). Thus, Piezo1 activation is sufficient to drive chromatin associated reorganization under static conditions. To connect the chromatin structural alterations to functional outputs, we performed SMT of KLF2 in the presence of Yoda1 treatment under static conditions. However, Yoda1 alone did not significantly alter KLF2 mobility compared to DMSO control (Fig. 5J-K) despite a trend toward FSS-induced confinement (Fig. 2 C-D). This is consistent with the known inactivation kinetics of Piezo1, while Yoda1 slows the inactivation of mechanically evoked currents [25]. Under continuous FSS, the channel is repeatedly activated by sustained mechanical stimulation, maintaining elevated Ca²⁺ signaling across the full 6 hr window during which SMT was measured. In contrast, prolonged exposure may result in Piezo1 signaling feedback including calcium signaling loss and receptor uncoupling [26].

**Figure 6:**
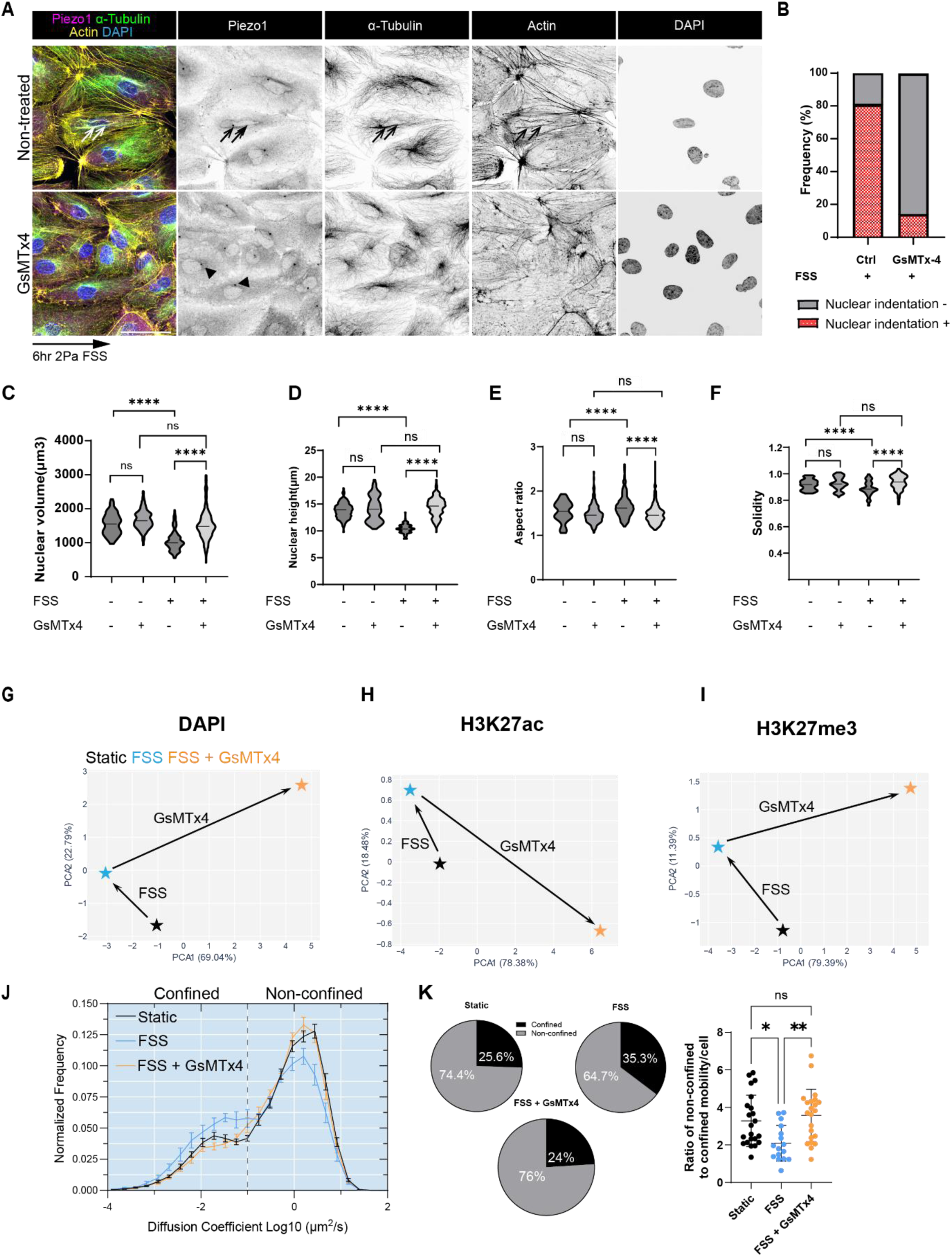
Piezo1 signaling plays a role in chromatin organization and TF mobility. (A) Representative MIPs of HAECs exposed to FSS for 6hr in the absence or presence of the Piezo1 inhibitor, GsMTx4. Cells were stained for Piezo1 (magenta), α-Tubulin (green), F-actin (yellow) and nuclei (DAPI, blue). Arrows indicate Piezo1 localization at perinuclear linear cytoskeletal bundles after 6h FSS exposure. Arrowheads indicate perinuclear localization of Piezo1 in control cells. Scale bar, 50 μm. (B) Percentage of HAECs with at least one nuclear indentation under control and GsMTx4-treated conditions during FSS exposure. (n=100 cells from three independent experiments). (C-F) Quantitative analysis of nuclear morphology in HAECs exposed to FSS in the presence or absence of GsMTx4. Nuclear volume (C), height(D), aspect ratio(E) and solidity (F). (n=150 cells from three independent experiments). (G-I) PCA of features extracted from (G) DAPI, (H) H3K27ac, and I) H3K27me3 staining in HAECs under static conditions or exposed to 2 Pa FSS for 6 hrs with or without GsMTx4 (mechanosensitive channel inhibitor). Each star represents the condition centroid in PC space with arrows indicating the direction and magnitude of chromatin state change. Variance explained by PC1 and PC2 is indicated on each axis. (J) Mobility distribution plot comparing diffusion coefficients across static, FSS, and FSS + GsMTx4 conditions. Dotted line indicates the threshold between confined and non-confined mobility states. Data presented as mean ± SEM. (K) Pie charts represent the proportion of the population found in either confined or non-confined state based on diffusion coefficient. Dot plot represents the ratio of non-confined to confined molecules per cell, n ≥ 16 cells per condition from three biological replicates. Statistical significance was determined by Welch’s T-test, *p < 0.05, **p < 0.01.

Yoda1-stimulated signaling under static conditions is likely to have substantially diminished by this timepoint, generating a transient rather than sustained signal. While Piezo1 pharmacological activation was sufficient to reproduce several hallmarks of the FSS-induced nuclear mechanoresponse, it remained unclear whether broader mechanosensitive ion channel activity is required for these adaptations. We therefore inhibited mechanosensitive ion channels using GsMTx4 and assessed the consequences on flow-induced nuclear remodeling.

### Broad inhibition of mechano-sensing ion channels rescues FSS-induced nuclear remodelling and genome adaptive response

While Piezo1 pharmacological activation was sufficient to reproduce several hallmarks of FSS-induced nuclear mechanoresponse, it remained unclear whether broader mechanosensitive ion channel activity is required for these adaptations. We therefore inhibited mechanosensitive ion channels using GsMTx4 and assessed the consequences on flow-induced nuclear remodeling. Treatment with GsMTx4 in the presence of FSS significantly reduced Piezo1 localization at perinuclear cytoskeletal bundles compared to control cells (Fig. 6A, black arrows). Loss of these structures was accompanied by a pronounced reduction in the proportion of nuclear indentation-positive cells, with approximately 80% of control cells exhibiting nuclear indentations compared with only ∼15% of GsMTx4-treated cells (Fig. 6A-B). Consistent with the reduction in nuclear indentation, quantitative nuclear morphometric analysis further demonstrated that GsMTx4 treatment also prevent FSS-induced changes in nuclear architecture, including the reduction in nuclear volume, height, and solidity (Fig. 6C, D and F) and the increase in aspect ratio (Fig. 6E). Of note, under static conditions, GsMTx4 treatment did not significantly affect nuclear volume, height, aspect ratio, or solidity compared with untreated controls, further validating that mechano-sensitive ion channels are required for flow-induced adaptation. These findings suggest that beyond Piezo1, integrated signaling of mechanosensitive cation-channel is required for the formation or stabilization of perinuclear cytoskeletal bundles associated with nuclear deformation under flow.

We next investigated whether GsMTx4 treatment also restored chromatin associated organization toward the static state. MIEL analysis showed that GsMTx4-treated and FSS-stimulated cells occupied feature-space positions that were separated from both static and untreated FSS populations across DAPI, H3K27ac, and H3K27me3 channels (Fig. 6G–I, Supp. Fig. 6F–H). Consistent with this pattern, Euclidean distance analysis showed that GsMTx4-treated and FSS-stimulated cells were displaced from both static and untreated FSS conditions across all three channels (Supp. Fig. I–K). Spearman correlation analysis further showed that GsMTx4-treated and FSS-stimulated cells were negatively correlated with both static and untreated FSS feature states (Supp. Fig. 6L–N). Collectively, these analyses indicate that mechanosensitive ion channel inhibition does not restore chromatin organization to a pre-flow configuration. Rather, GsMTx4 uncouples flow-induced chromatin remodeling and drives the emergence of a distinct chromatin state that is molecularly different from both static and untreated FSS conditions.

Given that FSS was associated with both reduced nuclear volume and increased KLF2 confinement, we next explored whether these responses remained coupled to mechano-sensing when cation-channel signalling was inhibited. Under static conditions, GsMTx4 did not alter nuclear volume (Fig. 6C) and similarly had little effect on KLF2 mobility (Supp. Fig. 6O–P). During FSS, however, GsMTx4 shifted both readouts toward the static state, attenuating the FSS-associated decrease in nuclear volume and increase in chromatin-confined KLF2 (Fig. 6C, J–K). The coordinated rescue of nuclear volume and KLF2 mobility, despite the emergence of a distinct MIEL-defined chromatin state, suggests that transcription factor dynamics can remain coupled to nuclear morphology/volume while becoming uncoupled from broader epigenetic organization. This dissociation supports a model in which mechanosensitive regulation of nuclear morphology, including nuclear volume, contributes directly to the mobility state of KLF2, whereas chromatin organization shapes the genomic landscape available for KLF2 engagement without necessarily dictating its dynamic behavior.

Importantly, these findings do not propose a new physical principle linking nuclear architecture to transcription factor dynamics. Rather, they reveal how endothelial mechanotransduction exploits this principle to coordinate adaptive genome responses across biological scales. More broadly, our work provides a mechanistic framework linking force sensing at the plasma membrane to emergent genome dynamics and highlights nuclear morphology as a key intermediary through which mechanical forces influence gene regulatory programs.

## Discussion

Our findings establish a mechanistic framework linking force sensing at the plasma membrane to adaptive genome responses in endothelial cells. By integrating quantitative molecular imaging across scales, we show that Piezo1-dependent mechanotransduction coordinates cytoskeletal remodeling, nuclear architecture, chromatin organization and transcription factor dynamics during adaptation to fluid shear stress.

A major finding is the identification of transient nuclear indentation as an early endothelial response to FSS. Although endothelial nuclear flattening and elongation under flow have been described previously [27], it is unclear how early cytoskeletal remodeling physically reshapes the nucleus during the initial response to FSS. Our data reveal that shortly after flow onset (within minutes to hour), the perinuclear actin and microtubule cytoskeleton remodels from a mesh-like network into prominent apical linear cables that span the nucleus and locally deform the nuclear surface. This remodeling corresponds to the formation of a longitudinal groove on the apical nuclear membrane that aligns with the axis of endothelial polarization. These architectural changes coincide with reductions in nuclear volume and height, identifying nuclear indentation as an early structural intermediate linking cytoskeletal remodeling to nuclear adaptation during endothelial mechanotransduction.

The rapid appearance of these perinuclear apical linear cytoskeletal cables after FSS exposure is consistent with previous studies describing the perinuclear actin cap as a specialized force-transmitting cytoskeletal network mechanically coupled to the nucleus through the LINC complex required for nuclear rotation and reorientation [28, 29]. Together, our findings and previous studies support the notion that FSS-induced forces are transmitted from the cell surface to the nucleus through specialized perinuclear cytoskeletal architectures. In this framework, mechanotransduction operates through a continuous physical continuum linking plasma membrane force sensing to the nuclear interior.

An important implication of FSS-induced nuclear architectural changes is its potential impact on genome response. Previous studies have shown that apical actin fibers can deform the nucleus and are associated with local chromatin changes [30], while FSS has been shown to regulate endothelial chromatin condensation and histone modifications, including acetylation– and methylation-associated chromatin states [7]. Consistent with this, chromatin changes measured by MIEL and super resolution microscopy show that a sustained FSS induces time-dependent states transition via the remodelling of chromatin-associated feature rather than a single static shift from a baseline to a steady state flow equilibrium (Fig. 2E-G). DAPI derived texture features likely capture changes in global chromatin architecture associated with force-dependent nuclear deformation, whereas H3K27ac and H3K27me3 reports the organization of active and repressive marked regulatory regions. Together, the feature-space patterns observed across chromatin organization and histone marks distribution suggest that FSS engages related but mechanistically distinct layers of genome remodelling from meso– to nano-scale environments.

Notably, the chromatin texture and organization showed a tendency to return toward baseline following prolonged flow exposure, whereas endothelial cell morphology remained markedly altered, with cells retaining the characteristic elongated flow-aligned phenotype. This divergence suggests that cytoskeletal and cellular adaptation may operate on different timescales from chromatin adaptation. One possibility is that early chromatin remodeling functions as a rapid mechanosensitive response that primes transcriptional adaptation before being progressively refined and stabilized by longer-term regulatory programs. Further studies will be required to determine how these distinct layers of adaptation are coordinated and maintained during long-term exposure to flow. Collectively, these observations support a model in which endothelial adaptation to flow proceeds through a transient adaptive state rather than a direct transition between static and flow-adapted conditions. Shortly after flow onset, coordinated remodeling of nuclear architecture, chromatin organization and transcription factor dynamics generates a highly responsive intermediate state that differs from both static and long-term flow conditions. While several chromatin-associated features subsequently trend back toward baseline, persistent nanoscale remodeling and sustained endothelial alignment indicate that adaptation culminates in a distinct flow-adapted equilibrium. This framework suggests that endothelial mechanotransduction is intrinsically dynamic and involves temporally ordered responses operating across multiple biological scales.

In parallel to genome changes, we also observed a shift in KLF2 mobility toward increased confinement after 6hr of FSS, coinciding with reduced nuclear volume at the same timepoint. This co-occurrence is consistent with a physical model in which nuclear compression increases chromatin density within a smaller nuclear space, thereby increasing the probability of TF–chromatin encounters. Supporting this idea, SMT studies in developing zebrafish embryos found that reduced nuclear volume increases chromatin binding of structurally and functionally distinct transcription factors, including TBP and SOX19b [31]. Our findings extend this concept to a mechanosensitive endothelial context, suggesting that nuclear geometry may act as a physical regulator of TF–chromatin engagement. Based on this proof of concept it is tempting to speculate that not only KLF2 but more globally other TFs mobility more broadly will be impacted by changes in nuclear volume.

A surprising finding of this study was the dynamic redistribution of Piezo1 during the endothelial response to FSS. Under static conditions, Piezo1 redistributes to perinuclear force-transmitting structures.. Notably, pharmacological activation of Piezo1 with Yoda1 was sufficient to induce a similar redistribution, whereas Piezo1 inhibition prevented the formation of these structures, suggesting that Piezo1 activity and localization are closely linked during endothelial mechanoadaptation to continued FSS stimulation. Although Piezo channels are primarily found at the plasma membrane, they have also been reported at the nuclear envelope [32–34] where distinct channel populations have been implicated in nuclear regulatory processes, including control of meiotic checkpoint signaling. These observations raise the possibility that spatially distinct Piezo1 populations perform specialized signaling functions within the cell. Such an aspect of Piezo1 biology emerges as a previously underappreciated component of endothelial mechanotransduction.

Together, these findings support a model in which Piezo1-dependent cytoskeletal remodelling provides a mechanistic bridge between extracellular mechanical force and endothelial gene-regulatory adaptation, connecting a plasma membrane mechanosensor to the nuclear interior through a spatially and temporally ordered sequence of events. This study suggests that endothelial adaptation to flow is not driven solely by biochemical signaling cascades but instead emerges from the interplay between signaling pathways and dynamic changes in cellular architecture. In this model, progressive remodeling of nuclear volume and chromatin organization provides physical layers of regulation that shape transcription factor dynamics during adaptation to mechanical force.

## Materials and Methods

### Cell culture

Human aortic endothelial cells (HAECs) were purchased from Lonza (Lonza, CC-2535, lot-20TL231227) and used between passages 3-5. HAECs were cultured in EGM-2 (Lonza, CC-3024) supplemented with EGM-2 bullet kit (Lonza, CC-3124) containing 2% fetal bovine serum (FBS), bovine brain extracts, ascorbic acid, human epidermal growth factor, hydrocortisone, and antibiotics. Cells were maintained at 37°C in a humidified atmosphere containing 5% CO_2_.

### Transfection

For plasmid transfection, 1.2×10^5^ of HAECs were seeded and transfected at the same time using Lipofectamine ^TM^ 3000 transfection reagent (Thermo Fisher Scientific, L3000008), Opti-MEM and 1µg of plasmid as the manufacturer’s instructions. The medium was changed 4 hours after transfection and the cells were grown in the cell culture incubator for 24 hr and all the experiments were done 24hr post-transfection.

## In vitro stimulation of fluid shear stress

### Ibidi pump system

For application of shear stress, 1.2×10^5^ of HAECs were seeded in μ-Slide 0.4 Luer ibiTreat (ibidi 80176) precoated with fibronectin (Thermo Fisher Scientific, F0895). Before starting FSS experiments, EGM-2 supplemented with 2%FBS were substituted with the medium containing 10%FBS. Unidirectional laminar flow was applied to confluent monolayers using the ibidi pump system (ibidi, 10902) starting from 0.5Pa, 1Pa and 1.5Pa for 30minutes each and 2Pa for 6, 24 and 48hr respectively. Static monolayers used the same EGM-2 supplemented with 10% FBS and were cultured alongside flow-treated monolayers.

### Chemical Treatments

To inhibit mechanosensitive ion channels, Piezo1, cells were treated with GsMTx4 (Fujifilm, 4393-S) during exposure to fluid shear stress. GsMTx4 was added to the culture medium at a final concentration of 1 µM immediately prior to the onset of shear stress and maintained throughout 6 h FSS. Control cells were exposed to fluid shear stress in the absence of GsMTx4. Cells were treated with Yoda1 (1 µM, Tocris, 5586) under static culture conditions for 6 h. Control cells were treated with an equivalent concentration of DMSO for the same duration.

Taxol (10nM, ThermoFisher Scientific, P3456) and Nocodazole (100nM, ThermoFisher Scientific, 358240100), Latrunculin B (12.5nM, Merck Millipore, 428020) and control (DMSO) were applied to the endothelial monolayer and maintained throughout 6h FSS.

### Immunofluorescence and imaging

Cells were fixed with 4% PFA/PBS (15710, Electron Microscopy Sciences) at room temperature for 15 minutes, permeabilized with 0.1% Triton X-100 for 1hour and blocked with 1% bovine serum albumin at room temperature for 1hour. Primary antibodies were incubated at room temperature for 1hour in blocking buffer and secondary antibodies applied for 1 h at room temperature. Primary antibodies: mouse monoclonal antiPiezo1 (1:100, Novus Biologicals, NBP2-75617), rabbit monoclonal anti-alpha Tubulin (1:1000, Abcam, ab52866), rabbit monoclonal anti-gamma Tubulin (1:200, Abcam, Ab308382), rabbit monoclonal anti-VE cadherin (1:100, Invitrogen, 14-1449-82), rabbit monoclonal anti Lamin A/C [EPR4100] (1:200, ab108595, Abcam), rabbit polyclonal anti-Histone H3 (acetyl K27) (1:500, ab4729, Abcam) and mouse monoclonal anti-Histone H3 (tri methyl K27) (1:100, ab6002, Abcam). Secondary antibodies: anti-Mouse Alexa Fluor 647 conjugate (1:1000, Invitrogen, A21236), anti-Mouse Alexa Fluor 488 conjugate (1:1000, Invitrogen, A11008) and anti-rabbit Alexa Fluor 546 conjugate (1:1000, Invitrogen, A11035). For visualization of F-actin, Alexa Fluor Plus 647 Phalloidin (1:1000, ThermoFisher Scientific, A30107) or Alexa Fluor 546 Phalloidin (1:1000, ThermoFisher Scientific, A22283) was used together with secondary antibody staining. For nuclei labelling, 14 μM DAPI (ThermoFisher Scientific, 62248) was used. The cells were fixed with 4% PFA/PBS (15710, Electron Microscopy Sciences) at room temperature for 15 minutes.

Images were acquired using Olympus FV3000 confocal microscope with 20X/0.75 NA objective and UPlanXApo ×60/NA 1.42 oil immersion objective. Images were processed using Fiji software.

### Airyscan Super-Resolution Imaging

Representative Airyscan super-resolution images were acquired using a ZEISS LSM 980 microscope equipped with an Airyscan 2 detector in Multiplex 4Y mode using a 63×/1.4 NA oil immersion objective. Z-stacks were acquired at 0.13 μm intervals and processed using ZEN Blue Airyscan reconstructionn and analyzed with Fiji software for linescan analysis.

### Live cell imaging

For checking the dynamics of Piezo1 under FSS, HAECs were transfected with Piezo1-Halo Tag (Addgene plasmid 207834). Prior to the flow experiment, the cells were treated with Janelia Fluor 549-Halo Tag Ligand (0.5nM, Promega, GA1110) for 30minutes and gently washed 3 times. The attached u-slide is placed into the temperature-controlled stage top incubator (Ibidi, 12720) at 37°C for the stage. Using the ND multipoint acquisition, multiple cells were chosen to be imaged simultaneously. Images were taken at intervals of 3min for 2h at static condition and 6hr under FSS exposure with 100 ms exposure time and a 25% power of the 546 nm-laser line. Confocal z-stacks were acquired using an inverted Olympus IX83/Yokogawa CSU-W1 spinning disc confocal microscope equipped with a Zyla 4.2 CMOS camera (Andor).

### Super resolution imaging (dSTORM)

#### dSTORM image acquisition

dSTORM was performed on HAECs fixed under static conditions (0hr) or after 48hrs of FSS and immunolabelled for H3K27ac using an anti-H3K27ac primary antibody (Abcam, ab4729) and an Alexa Fluor 546-conjugated anti-rabbit secondary antibody (Invitrogen, A11035). Datasets were acquired on a Zeiss ELYRA PS.1 inverted microscope configured at a 61° incidence angle for highly inclined and laminated optical sheet (HILO) illumination. Cells were imaged using a 561 nm excitation laser through a 100×/1.46 NA oil-immersion TIRF objective. Emission was collected using a BP 570–620/LP 750 filter set and recorded on an iXon885 EM-CCD camera (Andor). Image acquisition was performed using Everspark 1.0 (Idylle).

#### Single-molecule localization and preprocessing

dSTORM acquisitions were analyzed in ThunderSTORM within ImageJ/Fiji. The first 1,000 frames were discarded to allow fluorophore equilibration. Localizations were detected using wavelet-based background subtraction filtering (B-spline order 3, σ = 1 and 4 pixels), followed by sub-pixel localization via iterative 2D Gaussian fitting [35]. Retained localizations were required to have a fitted PSF width of 100–300 nm, localization uncertainty <50 nm, and photon count between 300 and 10,000. Drift was corrected using ThunderSTORM redundant cross-correlation (RCC) with 1,000-frame segments [36]. Repeated detections of the same fluorophore were merged when localizations occurred within 30 nm and ≤5 consecutive dark frames [37]. Merged events containing >20 detections were excluded as likely artefacts. Final localization tables were exported as CSV files for downstream cluster analysis.

For each cell, the nuclear boundary was defined by a localization density mask computed on a 500nm grid and thresholded at the 15th percentile of non-zero bins. The nuclear interior was sampled using a non-overlapping grid of 2µm × 2µm regions of interest (ROIs). ROIs were accepted for analysis only when ≥90% of their area fell within the nuclear boundary.

#### Nanocluster identification and characterization

H3K27ac nanoclusters were identified within accepted ROIs using DBSCAN with ε = 60 nm and minPts = 3, selected from k-nearest-neighbor distance analysis of representative cells. Clusters with fewer than 5 localizations or convex hull area >500,000 nm² were excluded as noise or imaging artefacts. For each retained cluster, convex hull area, equivalent diameter, number of localizations, and circularity were calculated. Per-cell cluster density, mean cluster diameter, mean localizations per cluster, and clustered fraction were then calculated across accepted ROIs. Clustered fraction was defined as the proportion of localizations assigned to retained clusters, weighted by ROI localization count. These metrics are consistent with established approaches for quantifying H3K27ac nanocluster organization [38].

#### Spatial analysis of nanocluster distribution

To assess nanocluster spatial organization, Ripley’s L(r)−r was computed from cluster centroid coordinates pooled across accepted ROIs for each cell. Border-corrected Ripley’s K was estimated using the convex hull of centroid coordinates as the study region and transformed to L(r)−r, where zero indicates complete spatial randomness [39, 40]. Radii from 10–1000 nm were evaluated in 60 equally spaced steps. The CSR crossover radius was defined as the r value at which L(r)−r crossed zero and mean nearest-neighbor distance between cluster centroids was calculated across all nuclear clusters per cell.

### Single molecule tracking

#### Cell preparation

HAECs were transfected with Halo-tagged KLF2 (GeneCopoeia, EX-Z7737-M50). Prior to imaging, 1nM of Halotag-ligand (Promega, GA1110) was added directly to the imaging chamber for 10 minutes at 37C. Following incubation, cells were washed and imaging media composed of EGM with 10% FBS was added for FSS stimulation and imaging.

#### SMT acquisition

SMT movies were acquired on a Zeiss ELYRA PS.1 inverted microscope configured at a 61° incidence angle for HILO illumination. Cells were imaged using a 561 nm excitation laser through a 100×/1.46 NA oil-immersion TIRF objective. Emission was collected using a BP 570–620/LP 750 filter set and recorded on an iXon885 EM-CCD camera (Andor). To measure the diffusion coefficient of KLF2 a frame rate of 50 Hz (20ms exposure per frame) was used to acquire 6000 frames without intervals. To keep the cells under 2 Pa of pressure, the ibidi u-slide with connection tubes was put in a heated stage tabletop incubator (ibidi, 12720) and then attached to the ibidi flow system. 2 Pa of pressure was applied during all SMT acquisitions.

#### SMT analysis

Masking and segmentation of the nucleus was performed in ImageJ. To identify and reconstruct molecular trajectories, a custom-written MATLAB implementation of the multiple target tracing (MTT) algorithm, known as SLIMfast was used [41]. Parameters used for analysis were selected based on established literature [42–47]: localization error Localization error: 10^-6.5^, blinking (frames) = 2 (equivalent to on gap frame between detections), max # of competitors: 3, max expected diffusion coefficient = 3μm^2^/sec, box size = 7, timepoints = 8, clip factor = 4. The first four timepoints of each trajectory were used to calculate the mean squared displacement. Diffusion coefficient was calculated from each trajectories’ mean squared displacement and plotted. A value of 0.1μm^2^/s was used to define the boundary between confined and non-confined populations [47]. The mobility fractions of each cell were calculated by computing the number of trajectories whose mean diffusion coefficient was determined.

### Quantification of Nuclear Indentation–Positive Cells

Cells were classified nuclear indentation–positive if the apical nuclear surface contained at least one indentation associated with an overlying linear cytoskeletal bundle. The number of nuclear indentation–positive cells was manually counted from confocal images and normalized to the total number of cells in each field, determined by nuclear DAPI staining. Data are presented as the percentage of nuclear indentation–positive cells relative to the total cell population.

### Line Scan Analysis

To assess the spatial relationship between cytoskeletal structures, Piezo1 and nuclear morphology, fluorescence intensity profiles were obtained in Fiji/ImageJ using the Plot Profile function. A line was drawn across the apical nuclear indentation and the associated cytoskeletal bundle in Airyscan images. Fluorescence intensities for each channel were extracted along the same line and normalized to the maximum intensity of each channel. Normalized intensity profiles were plotted as a function of distance to visualize the spatial distribution and overlap of the respective signals.

### Nuclear morphometric analysis

Three-dimensional confocal image stacks were acquired with voxel dimensions of 0.24 × 0.24µm in x–y and 1.22 µm in z. Images of the nuclear channel were imported into Cellpose 4 and segmented in 3D mode, with anisotropy parameters adjusted to account for the axial resolution (z/x–y ratio = 5.1) and diameter 30 [48]. No additional training was performed. Following segmentation, nuclear masks were exported and analyzed using Fiji (ImageJ) and custom Python scripts. Nuclear volume was calculated by multiplying the number of voxels within each segmented nucleus by the voxel volume. Nuclear height was defined as the extent of the nucleus along the z-axis and calculated as the number of occupied z-slices multiplied by the z-step size (1.22 µm). Nuclear solidity was quantified in 3D as the ratio of the segmented nuclear volume to the volume of the corresponding convex hull, providing a measure of nuclear surface irregularity and deformation. Nuclear aspect ratio was determined from maximum intensity projections by fitting an ellipse to each segmented nucleus and calculating the ratio of the major to minor axis lengths.

All measurements were performed in physical units using native voxel dimensions, with anisotropic voxel scaling explicitly accounted for during segmentation and downstream analyses. Segmented objects were subjected to size and shape quality filters to exclude debris and segmentation artifacts.

### Image Processing and Feature Extraction for MIEL

Average intensity Z-projections were generated to reduce three-dimensional image stacks to two-dimensional representations. Cells were segmented from projected images using Cellpose v3.0 with the cyto3 model [49]. Image features were extracted from individual nuclei. To account for replicate-associated technical variation, feature values were adjusted using a linear regression model including experimental replicate as a covariate. Outliers were identified on a per-feature basis within each condition using standardized z-scores, with feature values having an absolute z-score ≥ 3 classified as outliers. For each cell, the fraction of features classified as outliers was calculated, and cells were excluded if more than 5% of measured features were flagged. Following outlier filtering, feature values were averaged within each biological replicate for each condition. PCA was then performed on the replicate-level feature matrix to visualize multivariate relationships between conditions, with the first two principal components plotted. Euclidean distances between condition centroids in PCA space were calculated to quantify relative displacement between conditions

#### Distance Metrics and Dispersion Analysis

Multivariate differences between conditions were assessed using PERMANOVA implemented in Python’s scikit-bio library (v0.6.3). Principal components were retained until they cumulatively explained >80% of total variance, and PERMANOVA was performed on Euclidean distances calculated across all retained PCs, with significance assessed using 999 permutations. Prior to PERMANOVA, homogeneity of multivariate dispersion was evaluated using PERMDISP with 999 permutations to assess whether group differences could be influenced by unequal within-group variability. When PERMDISP was non-significant, significant PERMANOVA results were interpreted as evidence of differences in group centroids within the retained multivariate PC feature space.

#### Spearman Correlation Matrix

Spearman correlation coefficients were calculated using the mean feature values aggregated from each replicate using Python’s scipy spearmanr library (version 1.16.0).

## Statistical analysis

Graphical presentations and statistical analysis was generated with Graphpad Prism Software v11.0 (Dotmatics). Details on individual plots are described in figure legends.

## Resource availability

Further information and requests for reagents may be directed to corresponding authors.

## Data and code availability

Datasets and source data can be found in DOI:10.5281/zenodo.20696393 and will be publicly available upon manuscript publication.

## Acknowledgements

We thank all members of the Phng lab and Francois lab for insightful discussions and valuable feedback. We also acknowledge RIKEN Kobe Bioimaging Facilities & Factory for providing advanced imaging systems and technical support.

## Author contributions

Conceptualization: H.W., M.G., L.K.P., and M.F., Performed experiments: H.W. and M.G. Resources: L.K.P. and M.F. Data analysis: H.W., M.G., and Y.W. Writing – original draft: H.W. and M.G. Writing – review and editing: H.W., M.G., L.K.P. and M.F. Supervision: L.K.P. and M.F. Project administration: L.K.P. and M.F.

---

**Supplementary Figure 1.**
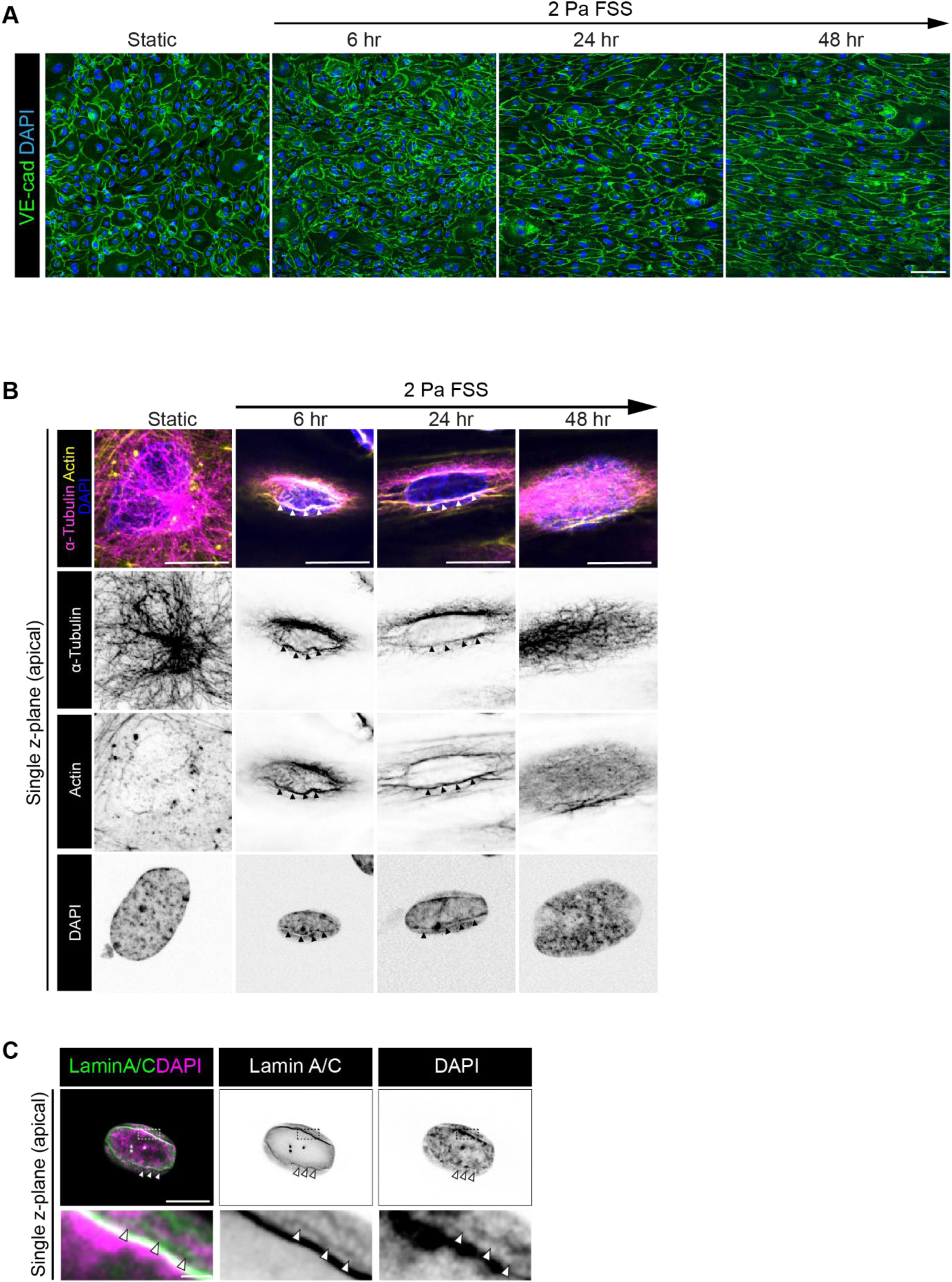
Laminar fluid shear stress induces endothelial alignment and transient nuclear indentation. (A) Representative low-magnification MIPs of HAECs cultured under static condition or exposed to laminar fluid shear stress (FSS, 2 Pa) for 6, 24 and 48 hr. Cells were stained for VE-cadherin (green) and nuclei (DAPI, blue). Scale bar,100 μm. (B) Representative single apical plane of the HAECs showing cytoskeletal and nuclear remodeling in response to FSS. HAECs exposed to 2 Pa FSS were stained for α-tubulin (magenta), F-actin (yellow) and nuclei (DAPI, blue). Arrow heads apical indentation of the nucleus at 6 and 24 hr after FSS exposure. Arrow indicates the direction and duration of FSS exposure. Scale bar, 20 μm. (C) Representative Airyscan image of a single apical plane of HAECs showing apical nuclear indentation 6hr after FSS exposure. Cells were stained for LaminA/C (green) and DAPI (magenta). Dashed box indicates apical nuclear indentation. White arrowheads, nuclear indentation with stripe of low DAPI signal on apical surface of nucleus. Enlarged images show regions highlighted by dashed boxes. Scale bars, 10 μm (upper panel) and 1μm (lower panel.

**Supplementary Figure 2.**
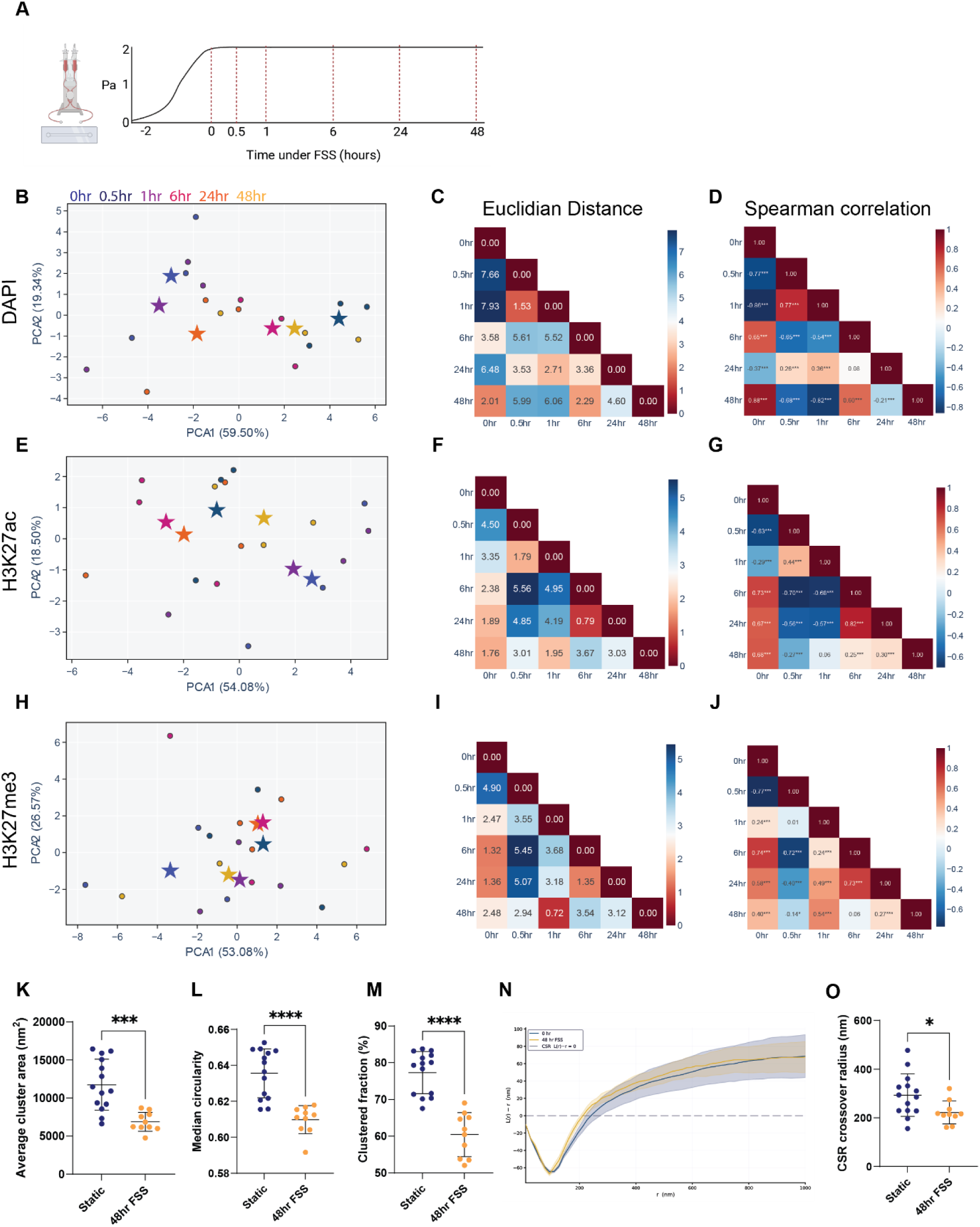
MIEL analysis of genome and chromatin organization across the FSS time course, and H3K27ac nanodomain characterization by dSTORM (A) Schematic of the FSS experimental timeline. HAECs were exposed to laminar FSS at 2 Pa and fixed at 0, 0.5, 1, 6, 24, and 48 hr for downstream analysis. (B–D) PCA of texture and edge-based features extracted from (B) DAPI, (C) H3K27ac, and (D) H3K27me3 staining in HAECs exposed to FSS for 0, 0.5, 1, 6, 24, and 48 hr. Each star represents the condition centroid and each dot an individual nucleus in PC space. Variance explained by PC1 and PC2 is indicated on each axis. Analysis performed on a minimum of n ≥ 750 nuclei per condition across n = 3 biological replicates. (E–G) Pairwise Euclidean distances between condition centroids in PC space for (E) DAPI, (F) H3K27ac, and (G) H3K27me3 channels. Larger values indicate greater dissimilarity in chromatin feature space between conditions. (H–J) Pairwise Spearman correlations of condition-level feature profiles across 252 features for (H) DAPI, (I) H3K27ac, and (J) H3K27me3 channels. Positive values indicate similar and negative values indicate opposing chromatin organization between conditions. Asterisks denote statistical significance (***p < 0.001). DAPI, H3K27ac, and H3K27me3 channels were analysed independently as each reflects a distinct dimension of nuclear organization, capturing chromatin compaction, active enhancer mark distribution, and repressive mark distribution, respectively. (K–O) dSTORM-based quantification of H3K27ac nanoscale organization under static conditions and after 48hr FSS. Unless otherwise indicated, each point represents one cell, calculated from the mean or median value across all valid interior ROIs for that cell. Static n = 14 cells and 48 hr FSS n = 10 cells from two biological replicates. Data are presented as mean ± SD. Statistical significance for dot-plot panels was determined by Welch’s t-test. (K) Mean H3K27ac nanocluster area per cell. (L) Mean H3K27ac nanocluster circularity per cell, where 1.0 indicates a perfect circle and lower values reflect more elongated nanocluster morphology. (M) Clustered fraction per cell, representing the proportion of H3K27ac localizations residing within identified nanoclusters. (N) Ripley’s L(r)−r analysis of H3K27ac nanocluster centroid organization. Curves represent the per-cell mean averaged across valid ROIs, with shaded bands representing ± SD. The dashed horizontal line indicates complete spatial randomness. (O) CSR crossover radius, quantifying the spatial scale at which H3K27ac nanocluster positions transition from more regularly spaced than complete spatial randomness to more clustered than complete spatial randomness.

**Supplementary Figure 3.**
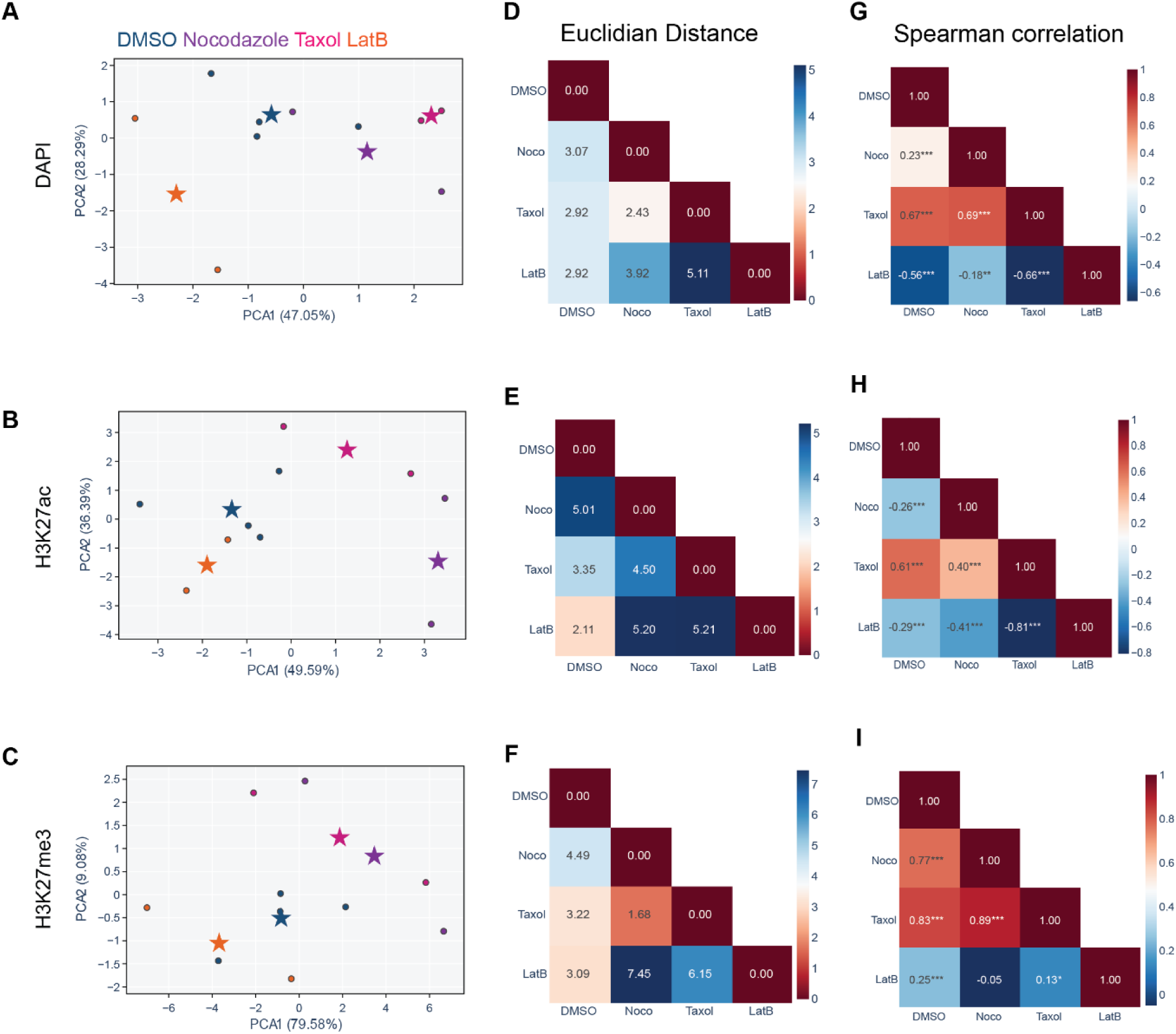
MIEL analysis of chromatin organization following cytoskeletal perturbation. (A-C) PCA of TAS features extracted from (A) DAPI, (B) H3K27ac, and (C) H3K27me3 staining in HAECs exposed to FSS for 6 hrs followed by treatment with DMSO (vehicle control), nocodazole (microtubule depolymerization), taxol (microtubule stabilization), or latrunculin B (Lat. B; actin depolymerization). Each star represents the condition centroid and each dot an individual biological replicate in PC space. (D-F) Pairwise Euclidean distances between condition centroids are shown for (D) DAPI, (E) H3K27ac, and (F) H3K27me3 channels, with larger values indicating greater dissimilarity in chromatin feature space. (G-I) Pairwise Spearman correlations of condition-level feature profiles are shown for (G) DAPI, (H) H3K27ac, and (I) H3K27me3 channels, with positive values indicating similar and negative values indicating opposing chromatin organization between conditions. Asterisks denote statistical significance (***p < 0.001, *p < 0.05). Analysis was performed on a minimum of 316 nuclei per condition, with 4 biological replicates for DMSO and 2 biological replicates per treated condition. Variance explained by PC1 and PC2 is indicated on each axis.

**Supplementary Figure 4.**
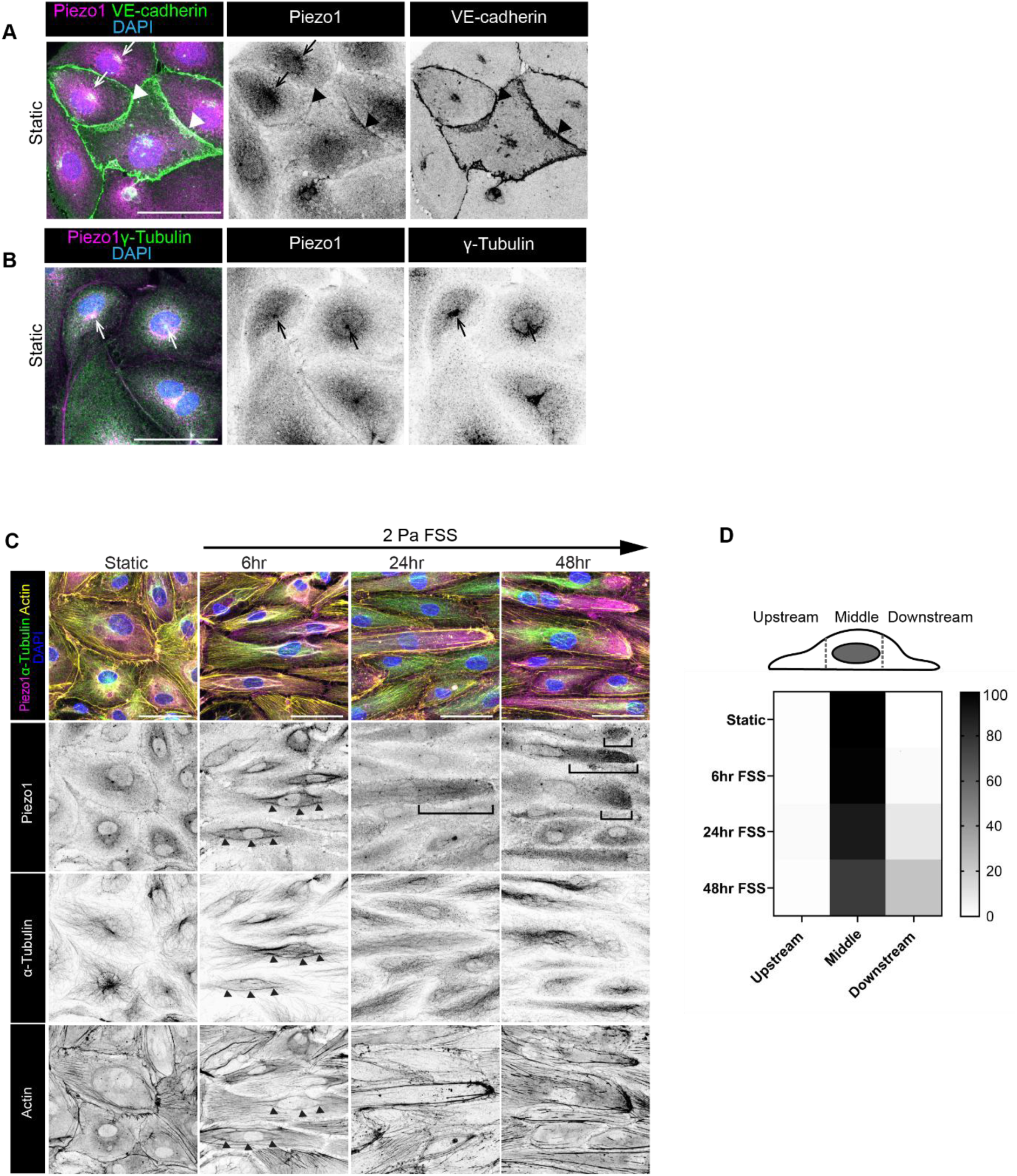
Piezo1 dynamically relocates during endothelial adaptation to fluid shear stress. (A) Representative MIPs of HAECs under static conditions stained for Piezo1 (magenta), VE-cadherin (green) and nuclei (DAPI, blue). Arrows, perinuclear localization of Piezo1. Arrowheads, Piezo1 localization at VE-cadherin-positive cell-cell junction. Scale bar, 50 μm. (B) Representative MIPs of HAECs under static condition stained for Piezo1 (magenta), γ-Tubulin (green) and nuclei (DAPI, blue). Arrows, colocalization of Piezo1 and γ-Tubulin at centrins. Scale bar 50 μm. (C) Representative MIPs of HAECs cultured under static condition or exposed to fluid shear stress (FSS, 2 Pa) for 6, 24 and 48 hr. Cells were stained for Piezo1 (magenta), α-Tubulin (green), F-actin (yellow) and nuclei (DAPI, blue). Arrowheads, perinuclear Piezo1 colocalizes with cytoskeletal bundles 6 hr after FSS exposure. Brackets, polarized Piezo1 localization at the downstream end of cells after 24 & 48 hr of FSS. (D) Heatmap summarizing the spatial distribution of Piezo1 intensity across HAECs exposed to FSS. Relative Piezo1 enrichment was quantified across upstream, middle and downstream cellular regions under static and flow conditions, n=100 cells from three independent experiments.

**Supplementary Figure 5.**
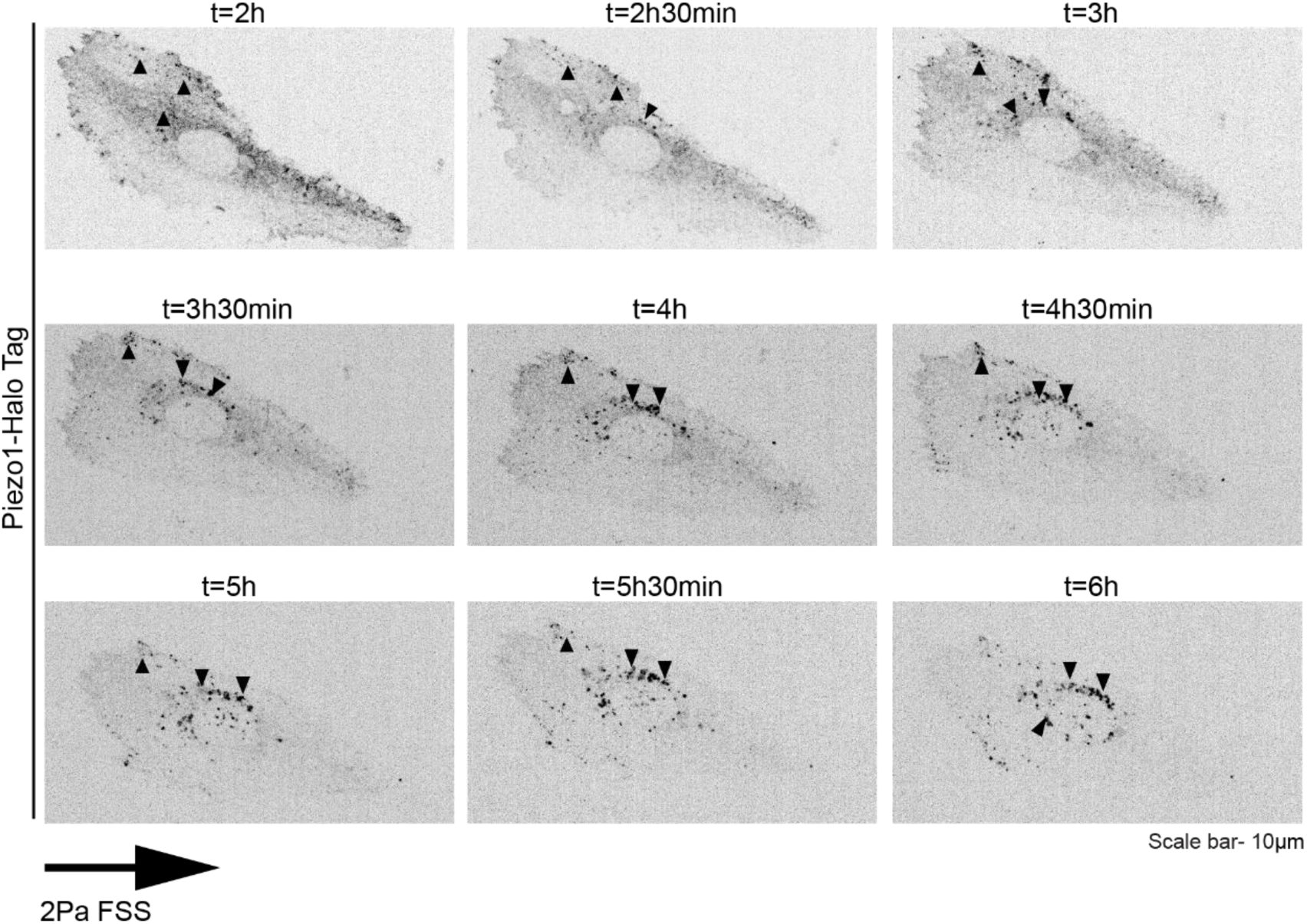
Time lapse images of live cell imaging of Piezo1-Halotag Frames are denoted by indicated time points. Arrows point to the temporal perinuclear localization of Piezo1 within 6hr FSS exposure. Scale bar, 10 μm.

**Supplementary Figure 6.**
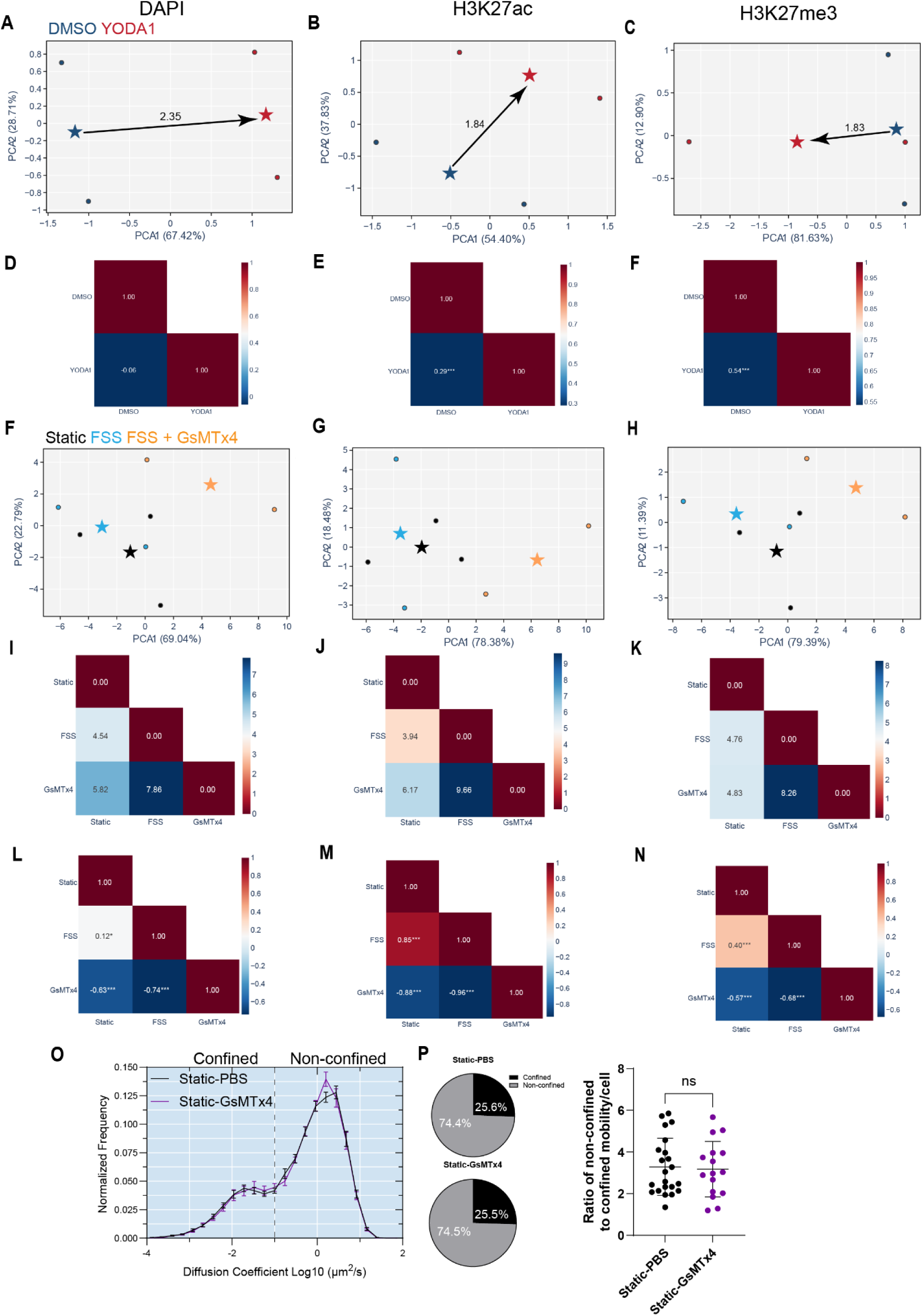
(A–C) PCA of features extracted from (A) DAPI, (B) H3K27ac, and (C) H3K27me3 staining in HAECs treated with DMSO (vehicle control) or YODA1 (Piezo1 agonist). Each star represents the condition centroid and each dot an individual biological replicate in PC space, with the Euclidean distance between centroids indicated. Variance explained by PC1 and PC2 is indicated on each axis. (D–F) Pairwise Spearman correlations of condition-level feature profiles for (D) DAPI, (E) H3K27ac, and (F) H3K27me3 channels between DMSO and YODA1 conditions. Asterisks denote statistical significance (***p < 0.001). (F–H) PCA features extracted from (F) DAPI, (G) H3K27ac, and (H) H3K27me3 staining in HAECs under static conditions or exposed to 2 Pa FSS for 6 hrs with or without GsMTx4 (mechanosensitive channel inhibitor). Each star represents the condition centroid and each dot an individual biological replicate in PC space. Variance explained by PC1 and PC2 is indicated on each axis. (I–K) Pairwise Euclidean distances between condition centroids for (I) DAPI, (J) H3K27ac, and (K) H3K27me3 channels, with larger values indicating greater dissimilarity in chromatin feature space. (L–N) Pairwise Spearman correlations of condition-level feature profiles for (L) DAPI, (M) H3K27ac, and (N) H3K27me3 channels. Positive values indicate similar and negative values indicate opposing chromatin organization between conditions. Asterisks denote statistical significance (***p < 0.001, *p < 0.05). (O) Mobility distribution plot comparing diffusion coefficients in HAECs under static conditions treated with GsMTx4. Dotted line indicates the threshold between confined and non-confined mobility states. Data presented as mean ± SEM. (P) Pie charts represent the proportion of the population found in either confined or non-confined state based on diffusion coefficient. Dot plot represents the ratio of non-confined to confined molecules per cell, n ≥ 16 cells per condition from three biological replicates. Statistical significance was determined by Welch’s T-test.

